# Bayesian phylodynamic inference of population dynamics with dormancy

**DOI:** 10.1101/2025.01.19.633741

**Authors:** Lorenzo Cappello, Wai Tung ‘Jack’ Lo, Joy Z. Zhang, Peiyu Xu, Daniel Barrow, Ishani Chopra, Andrew G. Clark, Martin T. Wells, Jaehee Kim

## Abstract

Many organisms employ reversible dormancy, or *seedbank*, in response to environmental fluctuations. This life-history strategy alters fundamental eco-evolutionary forces, leading to distinct patterns of genetic diversity. Two models of dormancy have been proposed based on the average duration of dormancy relative to coalescent timescales: weak seedbank, induced by scheduled seasonality (e.g., plants, invertebrates), and strong seedbank, where individuals stochastically switch between active and dormant states (e.g., bacteria, fungi). The weak seedbank coalescent is statistically equivalent to the Kingman coalescent with a scaled mutation rate, allowing the use of existing inference methods. In contrast, the strong seedbank coalescent differs fundamentally, as only active lineages can coalesce, while dormant lineages cannot. Additionally, dormant individuals typically mutate at a slower rate than active ones. Consequently, despite the significant role of dormancy in the eco-evolutionary dynamics of many organisms, no methods currently exist for inferring population dynamics involving dormancy and associated parameters. We present a Bayesian framework for jointly inferring a latent genealogy, seedbank parameters, and evolutionary parameters from molecular sequence data under the strong seedbank coalescent. We derive the exact probability density of genealogies sampled under the strong seedbank coalescent, characterize the corresponding likelihood function, and present efficient computational algorithms for its evaluation based on our theoretical framework. We develop a tailored Markov chain Monte Carlo sampler and implement our inference framework as a package SeedbankTree within BEAST2. Our work provides both a theoretical foundation and practical inference framework for studying the population genetic and genealogical impacts of dormancy.

## 1 Introduction

Many organisms respond to natural and anthropogenic disturbances by entering dormancy (*seedbank*), a reversible state of reduced metabolic activity [1–6]. This bet-hedging strategy enables populations to persist through unfavorable conditions, maintaining genetic, phenotypic, and functional diversity over time, thus serving as a temporal buffer against environmental fluctuations [5]. Dormancy fundamentally alters eco-evolutionary dynamics by generating age-structured populations with overlapping generations, thus affecting patterns of genetic diversity and population demography, even under neutrality [7–11]. This phenomenon is widespread across taxa, ranging from microbes to plants. For example, in microbial communities, over 90% of biomass in soils and marine sediments is estimated to exist in a dormant state [10, 12], illustrating the prevalence and ecological significance of seedbanks. Dormancy also has significant implications for infectious diseases. Pathogens, such as *Mycobacterium tuberculosis* (*Mtb*), can remain latent for years within hosts, evading immune responses and contributing to the long-term epidemiological burden of diseases [13, 14]. Such diverse and profound effects of the seedbank necessitate a fundamental understanding of dormancy as a key driver of eco-evolutionary dynamics.

Two fundamentally different models of dormancy have been proposed based on the average time individuals spend in the dormant state in comparison to the evolutionary timescale measured by the coalescent: the “weak” seedbank [7] that models dormancy induced by scheduled seasonality (e.g., plants or invertebrate species) and the “strong” seedbank [9, 15] where individuals stochastically switch between active and dormant states (e.g., bacteria or fungi). The weak seedbank model modifies the forward-in-time Wright-Fisher (WF) model to include seedbank effects, where individuals in each generation are drawn from seeds of multiple preceding generations according to age-dependent multinomial probabilities. This modification introduces age structure, reducing the effects of genetic drift and decreasing the rate of coalescence relative to the classical WF model, in which individuals are drawn only from the immediately preceding generation. The dual backward-in-time ancestral process converges weakly to the Kingman coalescent [16] with a reduced coalescence rate, reflecting the temporal dispersal of ancestral lineages due to the presence of the seedbank [7, 8]. Since the weak seedbank coalescent is equivalent to the Kingman coalescent with a population-rescaled mutation rate [11], existing statistical methods for the Kingman coalescent apply directly [17].

The strong seedbank model explicitly distinguishes between two populations: active and dormant. It arises as the backward-in-time dual of a WF model, where individuals stochastically transition between active and dormant states, with reproduction occurring exclusively among active individuals, while dormant individuals remain inactive for a geometrically distributed duration before reactivation. The limiting coalescent process of this WF model was first studied by Blath et al. [9, 15], who showed that the strong seedbank coalescent fundamentally differs from traditional coalescent models. While the presence of two populations resembles the two-population structured coalescent [18], a defining characteristic of the strong seedbank coalescent is that lineages in the dormant state cannot coalesce until reactivation, significantly extending coalescence times relative to standard coalescent models. The restriction that only active-state lineages can coalesce not only distinguishes the strong seedbank coalescent from the two-population structured coalescent, but also indicates that this process cannot be obtained through a simple time-rescaling of the Kingman coalescent.

These differences substantially modify genealogical structures compared to both the Kingman coalescent and the structured coalescent. Moreover, empirical evidence suggests that mutation spectra often differ between active and dormant states [5, 19], driven by reduced replication rates during dormancy [20, 21], differential oxidative damage [22–24], state-dependent DNA repair mechanisms [25, 26], and mutagenic stressors [27, 28] or adaptive mutagenesis [29, 30] associated with transitions between these states. These factors collectively reshape the mutational landscape and thereby the underlying evolutionary dynamics. The combined effects of genealogical and mutational processes generates distinct patterns of genetic diversity [9], which can be leveraged for statistical inference of dormancy and its associated parameters from molecular data [11]. This approach holds great promise given the eco-evolutionary significance of dormancy in many organisms. However, statistical inference under the strong seedbank model remains in its infancy [11]. Existing methods for other coalescent frameworks (e.g. [31–36]) are not directly applicable due to the distinct genealogical and evolutionary dynamics introduced by dormancy. Currently, there is no inference framework capable of jointly estimating seedbank population dynamics and its associated parameters, leaving a critical gap in the field. Our method addresses these challenges by introducing a novel, statistically rigorous computational framework for inferring population dynamics involving dormancy directly from molecular data without relying on summary statistics.

In this work, we develop, to our knowledge, the first Bayesian method for inferring eco-evolutionary dynamics under the strong seedbank coalescent, jointly estimating a genealogy, dormancy model parameters, and other relevant evolutionary model parameters from molecular data. Our method introduces several new components. We model the ancestral process with the strong seedbank coalescent, deriving a closed-form probability density for genealogies in bijection with this process, which serves as a prior for the latent genealogies representing the sample’s underlying evolutionary history. In addition, under general time-reversible substitution models, we formally characterize the the likelihood of molecular data given a genealogy, establishing an explicit connection to local-clock models (e.g., [37]). This formulation enables efficient phylogenetic likelihood computations using algorithms like Felsenstein’s pruning algorithm [38]. Furthermore, we develop a Markov chain Monte Carlo (MCMC) sampler to approximate the posterior distribution, leveraging recent methodological advances in structured coalescent-based inference. Lastly, we implement our method as the open-source package `SeedbankTree` within the `BEAST2` [39] platform and validate its accuracy and computational performance. We demonstrate its utility in uncovering insights into eco-evolutionary dynamics of dormancy using *Mtb* molecular data.

## 2 Background

Coalescent processes are stochastic models that define probability distributions on genealogies—timed, rooted binary trees representing the ancestral relationships within a sample from a population [40]. Their widespread utility stems from their robustness to underlying modeling assumptions, as diverse population models converge to the same coalescent process under appropriate rescaling of time. For example, both the WF and Moran models converge to the Kingman coalescent under classical neutrality assumptions [16].

The strong seedbank coalescent [15], hereafter referred to as the seedbank coalescent, models a population with a seedbank component, where lineages stochastically transition between active and dormant states. This model arises as the scaling limit of a WF model in which individuals can enter a seedbank and remain dormant without mortality during dormancy for geometrically distributed durations, or equivalently, from a Moran model incorporating seedbanks. The dynamics of the seedbank coalescent are characterized by three possible transitions: coalescence of two active lineages, transition of an active lineage into dormancy, and reactivation of a dormant lineage. Below, we formally define the seedbank *k*-coalescent.

### Definition 1 (The seedbank *k*-coalescent [15])

For a given sample size *k* ≥ 1, let 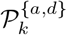 denote the space of “typed” partitions of [*k*], where each block of a partition is labeled as either active (*a*) or dormant (*d*). The *seedbank k-coalescent* is a continuous-time Markov chain (CTMC) on 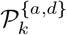, with transitions between *π*, 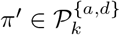 defined by:

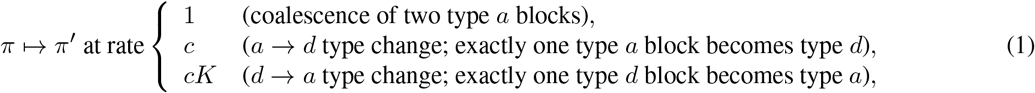

where *c* > 0 is the rate at which an active block becomes dormant, and *K* > 0 represents the relative size of the active population compared to the dormant population under the assumption of a constant population size.

A full realization of the seedbank coalescent is in bijection with a colored, timed genealogy, where the coloring indicates whether each lineage is active or dormant at a given time (Figure 1). We refer to a sample from the seedbank coalescent as a *seedbank genealogy*, which we formally define in the next section, along with the analytic formula for the probability distribution it induces on the associated genealogical space. The symbols and their definitions used in this work are summarized in Table S1.

**Figure 1.**
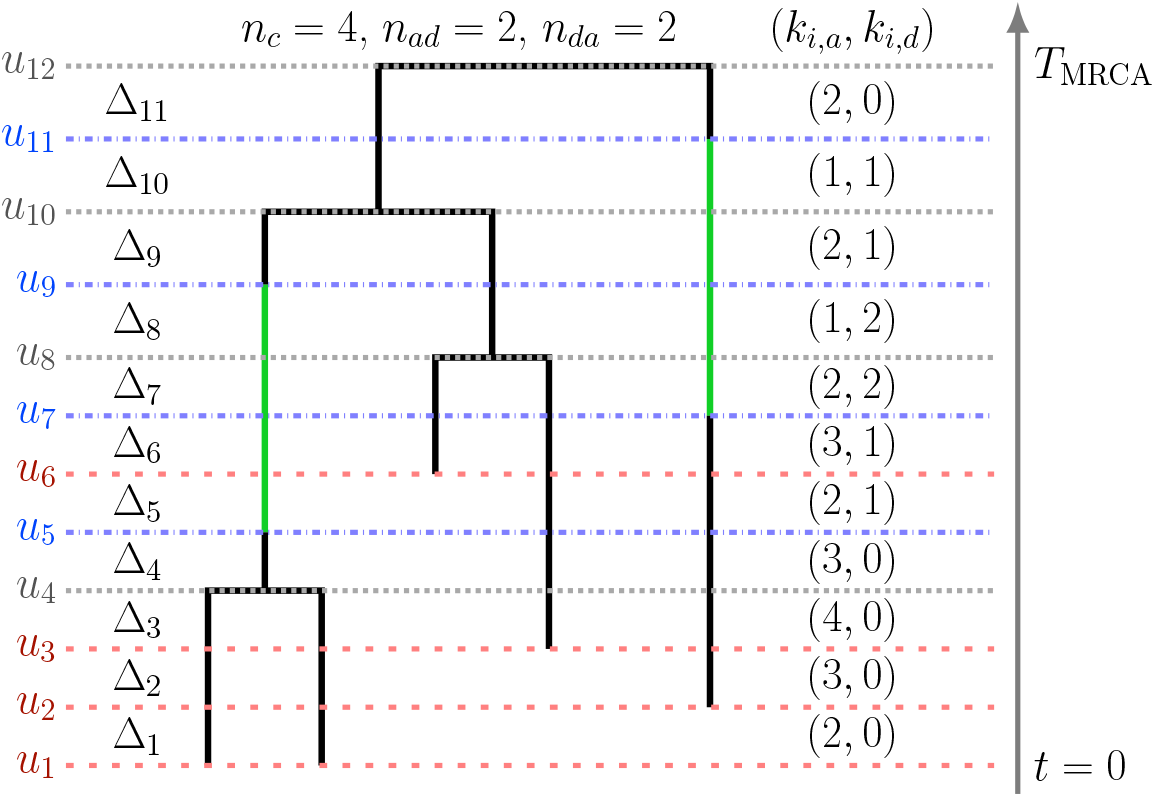
Example of a seedbank genealogy with sequential sampling. Active and dormant states are represented by black and green lineages, respectively. Dotted horizontal lines in red, gray, and blue illustrate sampling, coalescent, and type-change events (*a* → d or d → a), respectively. Each of *u*_1_, …, *u*_12_ denotes the time at which an event—sampling, coalescence, or type change—occurs, with times ordered increasing backwards in time (rootward) starting from the most recent sampling event at *u*_1_ = 0. *u*_12_ = *t*_MRCA_ indicates the time to the most recent common ancestor at the root. The time interval length between successive events is given by Δ_*i*_ = *u*_*i*+1 −_ *u*_*i*_ for the *i*-th interval (*u*_*i*_, *u*_*i*+1_). The pair (*k*_*i,a*_, *k*_*i,d*_) denotes the number of active and dormant lineages within this interval. The total counts of coalescent events, *a* →*d* type-change events, and *d* → *a* type-change events in the tree are respectively denoted by *n*_*c*_, *n*_*ad*_, and *n*_*da*_. For each edge *e*_*i*_ ∈*E*, the coloring of the edge (type-change events and their corresponding times) is defined by 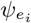.

Unlike classical coalescent models, such as the Kingman coalescent and the structured coalescent [18], which permit all lineages to coalesce within their respective demes, the seedbank coalescent imposes a key structural constraint: dormant lineages are excluded from coalescent events until reactivation. This restriction significantly impacts genealogical structure, including extending the expected time to the most recent common ancestor (*T*_MRCA_) [9, 11, 15]. Closed-form expressions for summary statistics of key genealogical properties, such as the first and second moments of *T*_MRCA_, are available [9]. For completeness, we provide comparative summaries of these theoretical distinctions between the structured coalescent and the seedbank coalescent (Table S2).

## 3 New Approaches

### 3.1 Genealogical model

#### Definition 2 (Seedbank genealogy)

We define a *seedbank genealogy* **g** = (*V, E*, **u**, *M*) with *n* samples as a ranked rooted binary tree augmented with a set *M* describing the type changes. The set *V* is the set of tree nodes partitioned into *V* = *S* ∪ *C*, where *S* is the set of *n* leaf nodes corresponding to sampling events, and *C* is the set of *n* − 1 internal nodes representing coalescent events between active lineages. We define *E* = {*e*1, …,*e*2*n*−2}to be the set of edges in **g**, defined by the nodes in *V* ; that is, the edges of a standard genealogy ignoring the type changes due to the transitions between active and dormant nodes. Let **u** = (*u*_1_, …, *u*_*B*_) be the vector of ordered event times in calendar-time units, each corresponding to a sampling, coalescence, or type-change event. We index each *u*_*i*_ such that time increases into the past (rootward) from the most recent sampling time (*u*_1_ = 0) to the time of the most recent common ancestor (*u*_*B*_ = *t*_MRCA_), the final coalescent event. Finally, 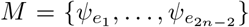 is a set of sample paths of type-change processes, with 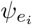 describing the transitions from active to dormant and vice versa (*a* → *d* or *d* → *a*) occurring on a given edge *e*_*i*_ ∈ *E*, along with their event times.

We denote the total number of coalescent events by *n*_*c*_ = |*C*| = *n* − 1. Given *M*, we can obtain the total counts of type-change events from active to dormant (*a* → *d*) and from dormant to active (*d* → *a*) along **g**, which we denote as *n*_*ad*_ and *n*_*da*_, respectively. Throughout the paper, we will informally refer to *M* as the set of coloring functions, as they represent the colors on the genealogy describing type-change events. We will also refer to the dormancy periods along an edge as colored portions.

#### Definition 3 (Reduced genealogy)

We define a *reduced genealogy* 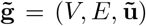 as a ranked rooted binary tree obtained from the seedbank genealogy **g** = (*V, E*, **u**, *M*) by removing *M*. The vector of event times **ũ** is derived from **u** by excluding the times corresponding to type-change events, thus retaining only the times of sampling and coalescent events.

An example of a seedbank genealogy with five active samples is shown in Figure 1, and the corresponding reduced genealogy is presented in Figure S2. Our definition of the seedbank genealogy closely follows that of the “structured tree” introduced by Vaughan et al. [33] for the structured coalescent. However, we consider only two types (active and dormant), and coalescent nodes are constrained to be of the active type, reflecting that coalescence occurs exclusively among active lineages.

Based on the formal definition of the seedbank genealogy, we derive the probability density of a seedbank genealogy under the seedbank coalescent model.

#### Theorem 1 (Probability density of a seedbank genealogy)

*Consider a seedbank genealogy* **g** *with n samples. Let θ* = *N*_*a*_*τ*_*g*_ *denote the scaling factor converting coalescent time units to calendar time, where N*_*a*_ *is the effective population size of the active population, and τ*_*g*_ *is the generation time in calendar units. For* 1 ≤ *i < B, define* Δ_*i*_ = *u*_*i*+1_ −*u*_*i*_ *as the length of the i-th time interval* (*u*_*i*_, *u*_*i*+1_). *Let k*_*i,a*_ *and k*_*i,d*_ *denote the numbers of active and dormant lineages, respectively, during this interval. Then, the probability density of a seedbank genealogy* **g** *under the seedbank coalescent is given by:*

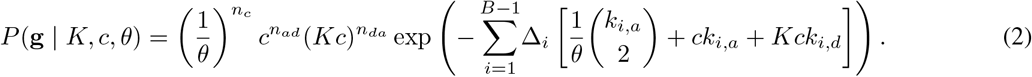

The proof follows directly from the properties of the CTMC seedbank coalescent process (Definition 1). The probability density is computed as the product of the contributions from the exponential waiting times between events and the instantaneous rates of events at their occurrence times. Eq. 2 is the product of holding times of the CTMC between events occurring within each time interval (*u*_*i*_, *u*_*i*+1_) of length Δ_*i*_, accounting for the cumulative rates of all possible events during that interval. At the end of each interval (*u*_*i*_, *u*_*i*+1_), exactly one event occurs at time *u*_*i*+1_—a coalescence between active lineages, an *a* → d type change, a *d* → a type change, or sampling—with the corresponding instantaneous rates incorporated into the overall probability density.

### 3.2 Mutation model

We employ a time-reversible CTMC model of molecular evolution [41, 42], with substitution rate matrices defined as **Π**_***a***_ = *μ*_*a*_**Q** for the active state and **Π**_***d***_ = *μ*_*d*_**Q** for the dormant state. The base rate matrix **Q** is normalized such that the expected number of substitutions is one per site per unit time, consistent with standard conventions in `BEAST2` [39]. *μ*_*a*_ and *μ*_*d*_ are mutation rates, corresponding to the expected number of substitutions per site per unit calendar time in active and dormant populations, respectively. We parameterize the dormant-state mutation rate as *μ*_*d*_ = *αμ*_*a*_. Since dormant lineages generally exhibit lower mutation rates than active lineages [5], we constrain *α* ∈ [0, 1]. We assume that the base rate matrix **Q** is identical across active and dormant states, based on the assumption that the underlying molecular mechanisms governing the mutational biases (e.g., oxidative stress [21]) and nucleotide equilibrium frequencies do not differ significantly between metabolic states. Although the overall mutation rates (*μ*_*a*_ and *μ*_*d*_) may vary with generation time [43], and latent-state mutagenic processes [25, 27] could bias mutation frequencies, we assume that the structure of the substitution model remains invariant across states.

Let **Y** = {**D, s**_**t**_, **s**_**s**_} represent the observed data, where **D** is the set of aligned molecular sequences, **s**_**t**_ denotes the corresponding sampling times, and **s**_**s**_ specifies the sample type (*a* or *d*) for each sequence. The phylogenetic likelihood *P* (**Y** |**g, Π**_*a*_, **Π**_*d*_) is the probability of the sequence data **Y**, given the seedbank genealogy **g** and the substitution model parameters. Given *M*, each edge in *E* may be partitioned into multiple segments corresponding to active and dormant states, each with different mutation rates. A naive approach to compute the likelihood of the seedbank genealogy is to use Felsenstein’s pruning algorithm [38], treating each type-change node as an additional internal node. This approach necessitates *n*_*ad*_ + *n*_*da*_ additional recursion steps, thereby increasing computational complexity. In Theorem 2, we prove that this is unnecessary. To eliminate the need to explicitly model state transitions along each branch, we introduce an “effective substitution rate matrix” for each branch that integrates the effects of both active and dormant periods into a single time-homogeneous Markov process.

#### Lemma 1 (Branch-specific effective substitution rate matrix)

*For each edge e*_*i*_ ∈*E, let* 𝓁_*i*_ *denote its length, and let* λ_*i*_ ∈ [0, 1) *represent the fraction of* 𝓁_*i*_ *that is “colored”. Then, the transition probability matrix along edge e*_*i*_ *is given by:*

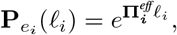

*where* 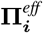 *is the branch-specific effective substitution rate matrix of edge e*_*i*_, *defined as:*

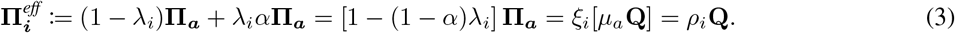

*Here, ξ*_*i*_ := 1 − (1 − *α*)λ_*i*_ ∈ (0, 1] *is the branch-specific scaling factor, and ρ*_*i*_ := *ξ*_*i*_*μ*_*a*_ *is the branch-rate multiplier, representing the expected number of substitutions per unit time along the edge e*_*i*_.

The branch-specific effective substitution rate matrix 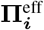encapsulates the combined effect of the active and dormant states weighted by the proportion of time λ_*i*_ spent in the dormant state along edge *e*_*i*_. By defining 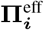, we effectively convert the heterogeneous substitution process into a homogeneous one with a modified rate matrix specific to each branch. Building upon this result, Theorem 2 shows that the phylogenetic likelihood of the seedbank genealogy **g** is equivalent to that of the reduced genealogy 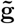, using branch-specific effective substitution rate matrices 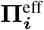.

#### Theorem 2 (Phylogenetic likelihood of the seedbank genealogy)

*The phylogenetic likelihood of a seedbank genealogy* **g** *is equal to that of its reduced genealogy* 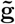 *with branch-specific effective substitution rate matrices* 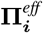 *for each edge e*_*i*_ ∈ *E, i* ∈ {1, …, 2*n* − 2}. *Consequently, the likelihood depends only on the substitution model parameters* {*α, μ*_*a*_, **Q**}, *edge lengths* 𝓁 = (𝓁_1_, …, 𝓁_2*n*−2_), *and dormant state proportions* **λ** = (λ_1_, …, λ_2*n*−2_).

The proofs of Lemma 1 and Theorem 2 are provided in Section S1 and Section S2 of the Supplemental Information, respectively.

Lemma 1 and Theorem 2 provide the theoretical basis for extending standard phylogenetic methods to the seedbank model, accounting for the impact of dormancy on molecular evolution and enabling efficient implementation using existing algorithms. By incorporating branch-specific effective rate matrices 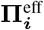 into the reduced genealogy 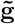, the phylogenetic likelihood of the seedbank genealogy **g** can indeed be computed using Felsenstein’s pruning algorithm [38] without additional computational cost. From a phylogenetic likelihood perspective, these branch-specific effective rate matrices, 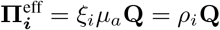, can be interpreted as the random local clock [37, 44, 45], where ***ρ*** = (*ρ*_1_, …., *ρ*_2*n*−2_) serves as the branch-specific rate multipliers.

### 3.3 Bayesian inference

We describe how to do Bayesian inference on the seedbank genealogy **g**, the seedbank parameters {*c, K, θ*}, and the mutation parameters {*α, μ*_*a*_, **Q**}. This framework is standard in coalescent-based phylodynamic inference. The target is the joint posterior density:

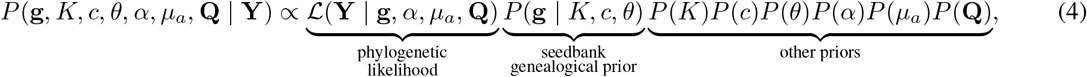

where the priors on *μ*_*a*_, **Q**, and *θ* follow from routine choices in phylodynamic inference (e.g., `BEAST2` [46]). The main distinction of Eq. 4 from existing literature lies in the prior on **g**, which is the seedbank coalescent (Theorem 1), the priors on the seedbank parameters, and the likelihood computation. The type-change rate *c* and the relative size of the active to dormant population *K* are positive real-valued parameters, hence we place a log-normal prior on them. The relative mutation rate between the active to dormant populations, *α*, is constrained within the interval [0, 1] through a Beta prior. Hyperparameters for the priors are chosen to control prior informativeness [47]. Details of the phylogenetic likelihood computation are provided in Section 3.2. Here, we consider the most general formulation: joint inference of mutation rates, population size, and population history, which requires serially sampled sequences [48].

The seedbank coalescent shares conceptual similarities with two classes of models: the structured coalescent (specifically, the two-deme coalescent [18]) and local clock models [49]. However, inference methods developed for these models are not directly applicable to the seedbank coalescent. Both the seedbank coalescent and structured coalescent utilize colored genealogies. In the structured coalescent, colors indicate the population assignments of lineages at a given time, with coalescence restricted to lineages within the same population (i.e., sharing the same color). Lineages may transition between populations over time through migration, altering their coloring. In the seedbank coalescent, coloring indicates whether a lineage is in an active or dormant state. Coalescent events are constrained to occur exclusively between active lineages, precluding dormant lineages from coalescing. Dormant states are represented on the edges of the genealogy and cannot appear at the nodes defining these edges, except at the tips when dormant sequences are directly sampled. An additional distinction lies in how state transitions influence the likelihood. Since mutation rates differ between active and dormant states, making changes in state assignments (coloring) directly impact the likelihood. This feature contrasts with the structured coalescent, where mutation rates are typically assumed to be identical across populations (colors), and changes in coloring do not influence the likelihood.

Similarly, local-clock models cannot be applied directly. Although both the seedbank model and local-clock models incorporate branch-specific rate multipliers ***ρ*** in likelihood computations, they differ fundamentally in how these multipliers are handled. In local-clock models, the rate multipliers are directly inferred, requiring careful consideration of their identifiability [37, 45]. In contrast, in the seedbank model, the rate multipliers are not directly inferred and do not appear explicitly in the posterior (Eq. 4). Instead, given a seedbank genealogy **g**, the coloring is used to compute **λ**, which, in combination with *α*, determines ***ρ*** via Eq. 3. Thus, the seedbank model avoids introducing 2*n* −2 additional parameters, as required in local-clock models, and reduces the parameterization to three key quantities: the transition rates governing active and dormant states (*c* and *K*), and the relative mutation rate *α*.

### 3.4 MCMC

We employ MCMC to approximate the posterior distribution (Eq. 4) by sampling from the target density. Efficient MCMC methods balance exploration and exploitation: new states are proposed to adequately explore high-density regions while avoiding chains getting stuck due to high rejection rates. Similar to `BEAST2`, we adopt a Gibbs-like strategy in which new states are proposed for a subset of variables, conditional on **Y** and the current states of all other variables not being updated. New states are proposed in blocks within a random-scan reversible MCMC framework [50]. The algorithm is initialized with a random sample from the prior distribution, after which state proposals are generated using a finite set of operators.

Most operators in our sampler use standard proposals, except those for the seedbank genealogy **g**, which require a custom-designed operator set tailored to the seedbank model (Section S3). We extend the structured tree operator design strategy of Vaughan et al. [33] to accommodate seedbank genealogies. Sampling a new seedbank genealogy involves two steps: (1) applying a tree operator to a reduced genealogy (Definition 3) of the current seedbank genealogy, and (2) proposing a new state assignment *M* for the subset of edges modified in step (1). The key in step (2) is to ensure that the proposed state assignment satisfies the seedbank model constraints. We detail this procedure next.

#### 3.4.1 Type-change proposal

Given an edge *e* ∈ *E* of length *l*, we propose a new state *ψ*_*e*_ by simulating an endpoint-conditioned CTMC from 0 to *l* over a finite state space. In the seedbank model, the CTMC consists of two states: active and dormant. By conditioning the chain on its start and end states, we enforce the constraints from Theorem 1. Specifically, if *e* = ⟨*υ, υ*^*′*^⟩ with *υ, υ*^*′*^∈ *C*, the CTMC is constrained to begin and end in the active state. Otherwise, if *e* = ⟨*υ* ❘, *υ*^*′*^⟩ with *υ* or *υ*^*′*^∈ *S*, the CTMC starts in the state from which it was collected and ends in the active state.

Several methods exist for sampling from an endpoint-conditioned CTMC with a finite state space (see Hobolth and Stone [51] for a comprehensive review). We employ the uniformization-based algorithm by Fearnhead and Sherlock [52]. Uniformization (e.g., [53]) enables sampling from any CTMC—not necessarily endpoint-conditioned—by introducing an auxiliary process. This auxiliary process comprises two components: a discrete-time Markov chain that determines the sequence of states visited by the chain and a Poisson process that specifies the number and timing of transitions. The scheme of Fearnhead and Sherlock [52] extends the standard uniformization by conditioning the auxiliary process on the endpoints, leveraging the simplicity of sampling from an endpoint-conditioned discrete-time Markov chain.

Let Ψ(*t*) be a CTMC with transition rate matrix **A**, whose off-diagonal entries represent active-to-dormant rate (*a*_*ad*_ = *c*) and dormant-to-active rate (*a*_*dc*_ = *cK*). Define *ν* = max {*c, cK*}, and construct a discrete-time Markov process Ψ^*d*^(*t*) with transition matrix 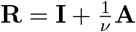 and a homogeneous Poisson process with intensity *ν*. By construction, **R** has one non-zero diagonal entry, indicating that Ψ^*d*^(*t*) permits self-loops. Hence, not all state changes sampled by the auxiliary process correspond to actual transitions. We refer to the sequence of states visited by Ψ^*d*^(*t*) as “virtual” events.

To sample a path Ψ(*t*) conditional on Ψ(0) = *x* and Ψ(*l*) = *y* (*x, y* ∈ {*a, d*}), we first sample the number of virtual state changes from the distribution:

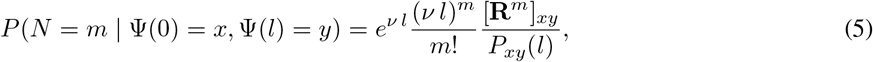

where *P*_*xy*_(*l*) := *P* (Ψ(*l*) = *y* | Ψ(0) = *x*). Intuitively, Eq. 5 modifies the standard Poisson process with rate parameter *νl* by incorporating endpoint conditioning. If *m* = 0, no state changes occur along the path. If *m* > 0, by the property of Poisson processes, we simulate *m* independent uniform random numbers from [0, *l*], sort them, and assign these times 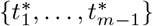 to the virtual-state changes in the path. Finally, we simulate 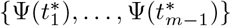 conditional on (Ψ(0) = *x*, Ψ(*l*) = *y*), as if simulating {Ψ^*d*^(1), …, Ψ^*d*^(*m* − 1)} given (Ψ^*d*^(0) = *x*, Ψ^*d*^(*m*) = *y*). This can be done with a forward-backward algorithm [54], which recursively defines the conditional transition distribution for *j* = 1, …, *m* − 1:

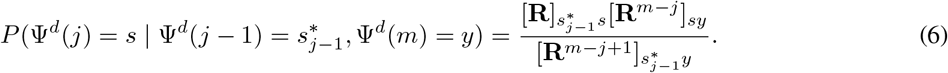

Given 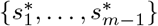, we obtain a sample from the endpoint-conditioned path of Ψ(*t*) by removing transitions that result in self-loops, along with the corresponding times. To summarize: (1) simulate the number of virtual events via Eq. 5, (2) simulate the times of the virtual events as uniform random numbers on [0, *l*], and (3) simulate the sequence of transitions using Eq. 6. For further details, see [52].

#### 3.4.2 Tree operators

We adapt the general tree operator strategy of Vaughan et al. [33] to the seedbank model, implementing a two-stage approach: (1) perform a genealogy move without considering branch coloring, and (2) propose a recoloring for all affected branches. The “uncolored” genealogy moves leverage the standard tree operators implemented in `BEAST2`, including the Wilson–Balding, subtree exchange, node-height shift, and tree-height scaling operators, which are widely used in coalescent-based inference [47, 48]. The recoloring step is specifically designed for the seedbank model, as detailed in Section 3.4.1, accounting for the constraints imposed by the active and dormant lineage states. The acceptance ratios for these operators are computed by combining the acceptance ratio of the uncolored genealogy move with that of the seedbank-specific recoloring proposal.

A key distinction between our seedbank model and the structured coalescent is that the coloring of the genealogy affects both the coalescent prior and the phylogenetic likelihood. As shown in Theorem 2, the effective mutation rate for an edge depends on the proportion of the edge in the dormant (colored) state, resulting in branch-specific effective rates analogous to those in local clock models. This dependency implies that, even with a fixed tree topology, variations in edge coloring are crucial for inferring edge lengths through coalescent times. To account for this, we introduce an “edge-recolor” operator that randomly selects an edge and proposes a new coloring without altering topology or branch length. While similar to the “node retype” operator of Vaughan et al. [33] and the “migration pair birth/death move” of Ewing et al. [32], our operator recolors the entire edge, rather than individual nodes or segments. The edge-recolor operator can add or remove dormancy along an edge, potentially altering the likelihood substantially.

Additional details of our tree operators are provided in Section S3 of Supplementary Information.

### 3.5 Implementation

We have implemented our method as the package `SeedbankTree` within `BEAST2`, enabling access to its comprehensive suite of substitution models, prior distributions, and MCMC operators. Our implementation extends the structured coalescent tree operators from the `MultiTypeTree` package developed by Vaughan et al. [33]. Our key advances include specialized data structures and computational routines tailored for integrating the seedbank coalescent model, along with seedbank-specific prior and likelihood computations. The seedbank tree data structure comprises explicit `SeedbankNode` nodes representing sampling and coalescent events, each annotated by its type (*a* or *d*). Implicit nodes for type-change events are stored within their immediate explicit child `SeedbankNode`, defining the edge in which they occur. To enforce the seedbank genealogy specification, we implemented a validation function that verifies consistency in node types and type-change events. We also implemented an initializer class, `SeedbankTreeInitializer`, that simulates a seedbank coalescent process to generate a random seedbank genealogy as an initial state of MCMC. `SeedbankTreeDensity` computes the probability density of a seedbank genealogy (Eq. 2) using the `SeedbankTree` data structure. The `TransitionModel` data structure manages the seedbank model parameters (*c, K*, and *θ*), and provides additional structures and helper functions to facilitate the uniformization procedure in the seedbank type-change proposal (Section 3.4.1). Finally, the seedbank branch rate model `SeedbankClockModel` computes the branch-specific scaling factors *ξ*_*i*_ (Lemma 1) for each edge by evaluating the proportion of each edge that is colored.

## 4 Results

We show the effectiveness of our methodology using both synthetic and real-world datasets. In the synthetic data analysis, we validate the computational framework through extensive simulations, demonstrating its accuracy by comparing analytically derived moments of summary statistics with MCMC-based estimates, as well as by recovering key seedbank coalescent and mutation model parameters across various parameter configurations and sampling schemes. In the real-data analysis, we apply the method to *Mtb* genomic data, inferring dormancy parameters, mutation rates, and genealogical structures that align with epidemiological observations.

### 4.1 Sampling from seedbank genealogical prior

To validate our implementation, we compared analytically derived first and second moments of key summary statistics, computed using recursive formulas from Blath et al. [9], with MCMC estimates drawn from the seedbank genealogical prior (Eq. 2). We focused on three statistics of the seedbank genealogy: the time to the most recent common ancestor (*T*_MRCA_) and the total lengths of active (*L*^(*a*)^) and dormant (*L*^(*d*)^) lineages. For *n* = 2, MCMC estimation was efficiently performed using either a single node-shift retype (NSR; Section S3.3) operator or a combination of tree scaler (TS; Section S3.2) and edge-recolor (RC; Section S3.6) operators. For larger sample sizes (*n* = 10 and 100), we employed the full operator set. The empirical means and variances of *T*_MRCA_, *L*^(*a*)^, and *L*^(*d*)^ closely matched their analytical counterparts (Tables S3–S5), confirming that our MCMC sampler accurately targets the seedbank genealogical prior.

### 4.2 Simulation study

We evaluated our inference method’s ability to recover underlying true model parameters using simulated data across four application-driven scenarios defined by (i) the type of data (molecular sequence vs. a known reduced genealogy) and (ii) the sampling scheme (isochronous vs. serial). Isochronous sampling collects all samples at a single time, whereas serial sampling spans multiple time points, enabling inference of time-resolved evolutionary dynamics [48]. We also considered sample types (active vs. dormant). Dormant samples are often unavailable in practice—for example, latent *Mtb* [23] and varicella zoster virus [55, 56] typically cannot be isolated from living hosts. However, spore-forming bacteria (e.g., *Bacillus subtilis* [57] or *Saccharomyces cerevisiae* [58]) and fungi (e.g., *Sordaria macrospora* [59]) could be sampled in the dormant state. Accordingly, we centered our analyses on two representative sampling scenarios: isochronous sampling with only active samples and serial sampling that includes both active and dormant samples. Because other sampling schemes—isochronous with both active and dormant or serial with only active (e.g., Section 4.3)—fall between these two scenarios, we focus on these two representative cases in this section.

Scenarios (1) and (2) use molecular sequence data to jointly estimate model parameters and the genealogy. They differ in their sampling schemes: isochronous in (1) and serial in (2). Scenarios (3) and (4) rely on a known reduced (uncolored) genealogy as observed data for model parameter inference under isochronous and serial sampling, respectively. This approach assumes the genealogy without coloring can be pre-estimated or known with reasonable accuracy, as commonly done in phylodynamic studies of both the Kingman coalescent [60–63] and the structured coalescent [34–36, 64].

Seedbank genealogies **g** were simulated under the seedbank coalescent. For scenarios (1) and (2), we further superimposed a mutation process on **g** using the HKY substitution model [65] with the transition/transversion bias *κ* to generate molecular sequences. For each parameter combination {*c, K, θ*} and {*α, μ*_*a*_ *κ*,} with a fixed sample size *n* = 25, we generated 100 replicates. Details of the simulation procedure are provided in Section 6.2. For each synthetic dataset, we approximated the posterior distribution by running an MCMC chain for 100 million iterations, discarding the initial burn-in, and thinning to obtain 20,000 posterior samples. Details on convergence assessment, burn-in selection, and prior hyperparameters are provided in Section 6.3.

To assess accuracy, we evaluated three metrics: coverage, relative bias, and relative root mean square error (RMSE). Coverage quantifies the proportion of replicates whose 95% posterior credible intervals include the true parameter value. Relative bias and relative RMSE are the bias and RMSE, respectively, scaled by the true parameter; for brevity, we simply refer to them as bias and RMSE:

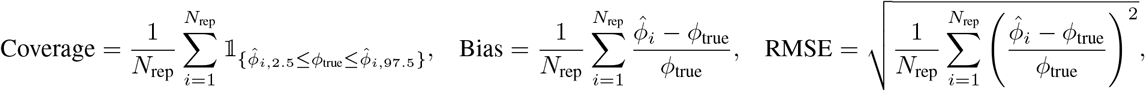

where *N*_rep_ is the number of replicates, 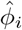 is the posterior mean from the *i*-th replicate, and *ϕ*_true_ is the true parameter value 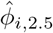 and 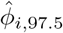 are the 2.5% and 97.5% posterior quantiles, respectively, for *ϕ* in the *i*-th replicate.

These metrics were computed for the three seedbank model parameters {*c, K, θ*} and the HKY parameter *κ* across all scenarios. Additionally, in scenarios (1) and (2) (molecular data), we included *T*_MRCA_, and in scenarios (2) and (4) (serial sampling), we evaluated the mutation rates *μ*_*a*_ and *α*. The results for scenarios (1) and (2) are presented in Table 1, while those for scenarios (3) and (4) are shown in Table S6. Both tables show that our method achieves near-perfect coverage for all parameters considered. Overall, accuracy is higher (i.e., lower bias and RMSE) in scenarios (3) and (4), which is expected because the genealogy is known, thereby eliminating genealogical uncertainty. The parameters specific to the seedbank coalescent (*c, K*) are estimated more accurately in the low-*K* regime across all scenarios. This improved accuracy can be partly attributed to the more balanced population sizes between active and dormant states, which results in a higher frequency of transitions between the two states.

**Table 1.** Coverage, relative bias, and relative RMSE of the posterior estimates for key model parameters and tree height, based on 100 replicate simulations, jointly inferring both the seedbank genealogy and the model parameters. Isochronous (Scenario 1; left) and serial (Scenario 2; right) sampling are shown. In isochronous sampling, all samples were active, whereas in serial sampling, samples included both active and dormant states. Reported parameters include the transition rate from active to dormant (*c*), the ratio of active to dormant population sizes (*K*), the scaled effective population size (*θ*), the time to the most recent common ancestor (*T*_MRCA_), the transition/transversion bias (*κ*), the dormant-to-active mutation rate ratio (*α*), and the active mutation rate (*μ*_*a*_). Columns indicate different true values of *K* and *α*. All other true parameter values were fixed at *c* = 1.0, *θ* = 1.0, *κ* = 3.0, and *μ*_*a*_ = 0.01. Under isochronous sampling, *α* and *μ*_*a*_ were not inferred, with results marked as not applicable (N/A). Results for serial sampling with *K* = 1 and *α* = 0.1 (marked “–”) are omitted due to inconsistent MCMC convergence across replicate runs.

|  | Isochronous sampling |  |  |  |  |  | Serial sampling |  |  |  |  |  |
| --- | --- | --- | --- | --- | --- | --- | --- | --- | --- | --- | --- | --- |
| | $K = 1$ | | | $K = 10$ | | | $K = 1$ | | | $K = 10$ | | |
| | $\alpha = 0.1$ | $\alpha = 0.5$ | $\alpha = 0.99$ | $\alpha = 0.1$ | $\alpha = 0.5$ | $\alpha = 0.99$ | $\alpha = 0.1$ | $\alpha = 0.5$ | $\alpha = 0.99$ | $\alpha = 0.1$ | $\alpha = 0.5$ | $\alpha = 0.99$ |
| Coverage |  |  |  |  |  |  | Coverage |  |  |  |  |  |
| $c$ | 1.00 | 1.00 | 1.00 | 1.00 | 1.00 | 1.00 | – | 0.99 | 0.99 | 0.99 | 1.00 | 0.99 |
| $K$ | 1.00 | 1.00 | 0.98 | 1.00 | 1.00 | 1.00 | – | 1.00 | 0.98 | 1.00 | 1.00 | 1.00 |
| $\theta$ | 0.94 | 0.97 | 0.96 | 0.99 | 0.97 | 0.94 | – | 1.00 | 0.99 | 0.94 | 0.97 | 0.99 |
| $T_{\text{MRCA}}$ | 0.97 | 0.95 | 0.95 | 0.97 | 0.93 | 0.97 | – | 0.95 | 0.93 | 0.90 | 0.95 | 0.94 |
| $\kappa$ | 0.94 | 0.98 | 0.95 | 0.94 | 0.93 | 0.96 | – | 0.98 | 0.97 | 0.97 | 0.96 | 0.96 |
| $\alpha$ | | N/A | | | N/A | | – | 0.97 | 0.98 | 1.00 | 1.00 | 0.99 |
| $\mu_a$ | | N/A | | | N/A | | – | 0.97 | 0.94 | 0.95 | 0.93 | 0.95 |
| Relative bias |  |  |  |  |  |  | Relative bias |  |  |  |  |  |
| $c$ | 0.042 | 0.12 | 0.12 | 0.12 | 0.12 | 0.14 | – | 0.088 | 0.12 | 0.025 | 0.14 | 0.13 |
| $K$ | 0.23 | 0.16 | 0.11 | 0.30 | 0.33 | 0.38 | – | 0.13 | 0.10 | 0.20 | 0.32 | 0.38 |
| $\theta$ | −0.016 | 0.039 | 0.039 | −0.011 | 0.041 | 0.020 | – | 0.068 | 0.14 | 0.048 | 0.076 | 0.056 |
| $T_{\text{MRCA}}$ | 0.034 | 0.0016 | 0.0020 | 0.0044 | −0.0094 | −0.0021 | – | 0.013 | 0.0066 | 0.0019 | 0.00055 | 0.00092 |
| $\kappa$ | 0.0060 | −0.00041 | −0.0024 | 0.0074 | 0.00042 | −0.000037 | – | 0.0016 | −0.0015 | 0.0030 | 0.0025 | 0.00039 |
| $\alpha$ | | N/A | | | N/A | | – | −0.031 | −0.047 | −0.0088 | −0.0041 | 0.089 |
| $\mu_a$ | | N/A | | | N/A | | – | −0.0058 | 0.021 | −0.0085 | −0.0046 | 0.012 |
| Relative RMSE |  |  |  |  |  |  | Relative RMSE |  |  |  |  |  |
| $c$ | 0.51 | 0.57 | 0.57 | 0.59 | 0.60 | 0.62 | – | 0.45 | 0.55 | 0.50 | 0.59 | 0.61 |
| $K$ | 0.66 | 0.58 | 0.54 | 0.74 | 0.78 | 0.83 | – | 0.44 | 0.45 | 0.55 | 0.72 | 0.82 |
| $\theta$ | 0.28 | 0.30 | 0.34 | 0.26 | 0.29 | 0.29 | – | 0.36 | 0.53 | 0.30 | 0.30 | 0.31 |
| $T_{\text{MRCA}}$ | 0.31 | 0.10 | 0.026 | 0.097 | 0.066 | 0.046 | – | 0.079 | 0.047 | 0.033 | 0.022 | 0.021 |
| $\kappa$ | 0.065 | 0.049 | 0.044 | 0.076 | 0.074 | 0.073 | – | 0.032 | 0.028 | 0.043 | 0.042 | 0.043 |
| $\alpha$ | | N/A | | | N/A | | – | 0.17 | 0.12 | 0.85 | 0.21 | 0.15 |
| $\mu_a$ | | N/A | | | N/A | | – | 0.065 | 0.042 | 0.054 | 0.046 | 0.034 |

The estimation accuracy of {*T*_MRCA_, *α, μ*_*a*_} is closely linked to the method’s ability to accurately reconstruct the true seedbank genealogy. While these parameters are accurately inferred, with nearly perfect coverage and small biases across scenarios, inference becomes more challenging as *K* and *α* decrease, resulting in larger absolute bias and RMSE. This is likely due to the inherent unidentifiability of the latent seedbank genealogy, as molecular data alone cannot fully resolve it. Bayesian inference can address this issue by constraining the parameter space via a prior distribution that assigns high probability to specific regions. Here, the seedbank coalescent, serving as the prior on the space of genealogies, provides such constraints only in certain parameter regimes.

Indeed, smaller values of *K* (i.e., a larger dormant population) increase seedbank genealogical uncertainty, with genealogies spanning a wide range of tree lengths and *T*_MRCA_ occupying high-probability regions (e.g., see Tables S3(B) and S5), thus complicating statistical inference. In contrast, for *K* = 10, seedbank genealogical uncertainty is substantially lower, leading to accurate inference. These issues become more pronounced when both mutation rates (*μ*_*a*_ and *α*) are also unknown; scenario (2) with *α* = 0.1, *K* = 1 is omitted from Table 1 because not all MCMC runs reliably converged due to poor mixing. A possible solution is to incorporate additional information, as demonstrated in scenario (4) with *α* = 0.1, *K* = 1 (Table S6), where all parameters are accurately inferred once the underlying uncolored genealogy is fixed.

Lastly, we replicated the simulation study using the standard Kingman coalescent as the prior for genealogies, effectively ignoring the seedbank effect. Applying this misspecified model to data generated under the seedbank coalescent enabled us to evaluate the bias introduced by this misspecification. Tables S7 and S8 present the results for {*T*_MRCA_, *μ*_*a*_, *κ*}, where the seedbank coalescent parameters are not inferred, and mutation rate *μ*_*a*_ is estimated only under serial sampling. With larger *K* and *α*, inference is relatively accurate, achieving near-perfect coverage when *α* = 0.99. However, as *α* and *K* decrease, coverage for {*T*_MRCA_, *μ*_*a*_} declines substantially, reaching zero in some scenarios. The *T*_MRCA_ is consistently negatively biased. This bias can be partially mitigated by either fixing the mutation rate to the effective mutation rate of the tree for isochronous samples (Table S7(A), right) or by directly estimating the mutation rate from the data for serial samples (Table S7(B)). In the latter case, while inference for *T*_MRCA_ improves, *μ*_*a*_ estimates remain inaccurate (bias, RMSE), with low coverage for both parameters. Incorporating additional information generally improves performance; however, overall inference remains substantially inaccurate (Table S8) compared to the correctly specified model (Table 1).

### 4.3 Application to *Mycobacterium tuberculosis* (*Mtb*)

Tuberculosis (TB) exhibits prolonged latency periods ranging from months to decades [66], with latent infections affecting approximately 25% of the global population [67]. Latently infected hosts are asymptomatic, making detection, sampling, and related parameter estimation particularly challenging [68]. We applied our inference framework to the whole-genome sequencing (WGS) dataset of *Mtb* from the outbreak of the Rangipo strain in New Zealand (1992–2011) [43]. Sampled individuals had established or putative epidemiological links, and some were hypothesized to have experienced prolonged latency. The authors reported significantly higher mutation rates in recently transmitted pairs than in latent pairs, with the latent-state rate at 0.13 times that of the active state.

The posterior mean estimates of seedbank and mutation model parameters align well with previous empirical findings from epidemiological studies. We inferred *c* at a posterior mean of 1.117 ([0.3667, 1.9946]), where the bracketed interval denotes the 95% HPD. Compared to previously reported doubling time in TB (95% CI: 0.64–1.82, corresponding to a transition rate from susceptible to exposed being 0.55–1.56 [69]), our estimate falls within the empirically supported range. The relative population size of active to dormant states *K* was estimated at 0.694 ([0.2509, 1.2636]). As *K* is not widely characterized in genetic epidemiology, we also sampled *cK*, the reactivation rate, obtaining a posterior mean of 0.786 ([0.1441, 1.6698]), corresponding to a latency period of 1.27 years, consistent with the reported median TB incubation period of 1.3 years from empirical studies [70]. The posterior mean of *κ* was 3.933 ([1.3519, 7.1212]), and the active mutation rate *μ*_*a*_ was 1.474 × 10^−7^ SNPs/site/year ([8.125 × 10^−8^, 2.1831 × 10^−7^]), close to the active mutation rate 2.409 × 10^−7^ reported originally for this dataset [43], and well within the TB mutation rate of 0.3–0.5 SNPs/genome/year range reported by Eldholm et al. [71]. The ratio of mutation rates between latent and active state, *α*, was estimated at 0.119 ([0.0476, 0.2003]), closely matching the reported value of 0.133 [43]. Finally, the mean *θ* was 5.345 ([0.8179, 12.0489]), and *T*_MRCA_ was estimated to be 1980.373 ([1968.608, 1989.765]). The maximum clade credibility (MCC) tree (Figure 2) also recovered the putative epidemiological links reported by Colangeli et al. [43] between samples *E* and *C*1, as well as between *S* and *O*.

**Figure 2.**
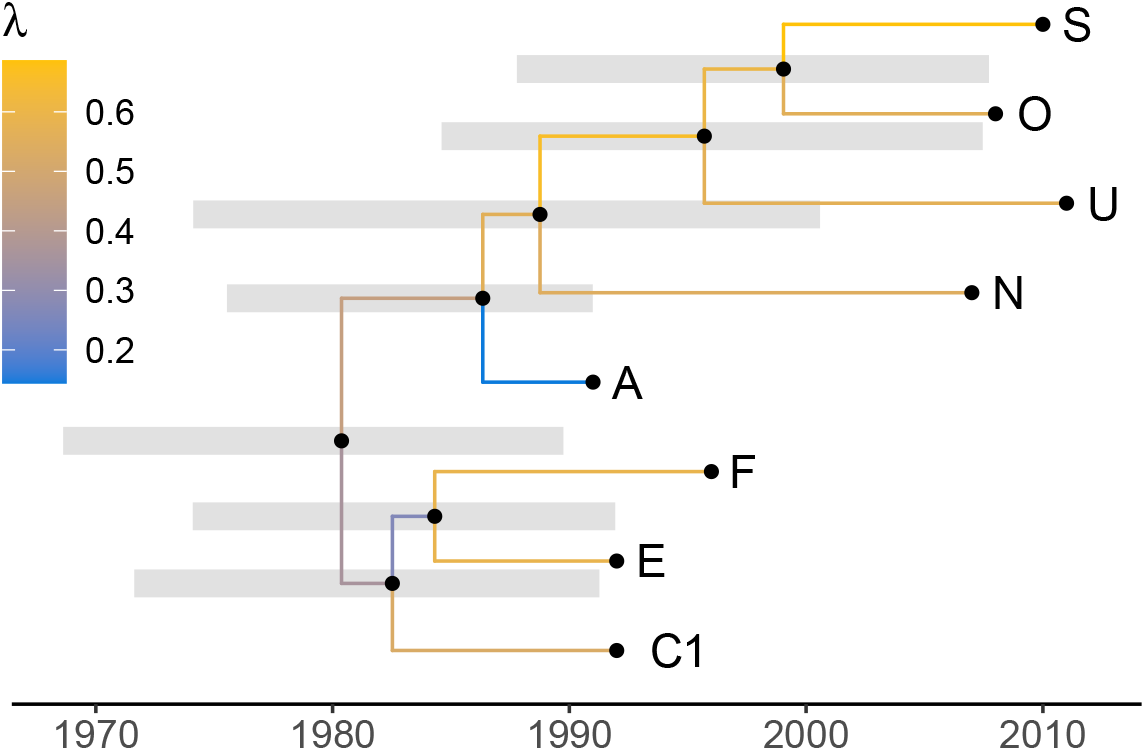
MCC tree of *Mtb* from the New Zealand dataset inferred using SeedbankTree. Branches are colored by the median proportion of latent state (λ) for each edge. All sampled tips are in the active state, and all internal nodes are also active, consistent with the seedbank model. Grey bars represent 95% HPD intervals for node heights.

## 5 Discussion

We have developed, to our knowledge, the first Bayesian method for jointly inferring a latent genealogy, dormancy parameters, and other relevant evolutionary parameters for populations experiencing dormancy. We derived the exact probability density of genealogies under the seedbank coalescent model, characterized the corresponding likelihood function, and developed a tailored MCMC sampler, implemented as an open-source package `SeedbankTree` within `BEAST2`. We also applied our method to *Mtb*, where latent-state physiology and mutation rate remain poorly understood, and metabolic-state mutation rate differences are often overlooked in phylodynamic studies due to the challenges of isolating latent sequences from living hosts. Our method provides both a theoretical foundation and practical inference framework for studying the population genetic and genealogical impacts of dormancy.

In both seedbank and structured coalescent models, genealogies are represented as colored trees where the color denotes distinct lineage states—e.g., active versus dormant lineages in the seedbank model or different subpopulations in the structured coalescent. The seedbank coalescent’s constraint that only active lineages can coalesce, while dormant lineages cannot, fundamentally alters the genealogical structure compared to the structured coalescent. Additionally, the structured coalescent typically assumes that the mutation process is independent of lineage state (color), such that the likelihood computation mirrors that of Kingman coalescent on an uncolored tree. However, in the seedbank model, this assumption no longer holds due to the differences in mutation rates between active and dormant states, necessitating the explicit incorporation of lineage state changes into the likelihood computation. This coupling under our model provides additional insights into population dynamics not captured by structured coalescent models.

To date, the only inference framework for the seedbank model is that of Blath et al. [11], which relies on the site frequency spectrum (SFS) as a summary statistic. While such SFS-based approaches are computationally efficient, their limitations for statistical inference are well-documented [72, 73]. Our work establishes the first foundational inference framework for the seedbank model from complete molecular data, thereby circumventing reliance on summary statistics. We provided numerical evidence identifying parameter regimes where accurate parameter inference is achievable. We further showed that ignoring the seedbank effect can bias estimates of key parameters, such as *T*_MRCA_ and the mutation rate. Additionally, we demonstrated that inference using molecular data alone becomes increasingly challenging as the dormant population’s mutation rate decreases and its size increases relative to the active population, likely due to inherent unidentifiability of the latent seedbank genealogy under these conditions. Incorporating additional data can mitigate this unidentifiability; we demonstrated one approach by fixing the uncolored genealogy. Further exploration of similar strategies remains a future research direction.

Our method provides a basis for future investigation of more complex demographic and environmental scenarios in diverse biological systems exhibiting dormancy. One natural extension is to relax the assumption of a constant population size, which is commonly done in phylodynamic inference using the Kingman coalescent [74–76] and has recently been explored within the structured coalescent framework [77]. Additionally, incorporating ancestral recombination graphs (ARGs) into the seedbank model would be an interesting extension, particularly given recent work on structured coalescent models with fixed ARGs [36]. More biologically realistic extensions would include a model in which multiple lineages undergo simultaneous transitions between active and dormant states in response to shared environmental pressures, as suggested by recent theoretical advancements [78]. Developing theoretical foundations that incorporate age-dependent reactivation and mortality rates during dormancy would also further enhance our understanding of dormancy’s impact on population dynamics. Lastly, integrating the seedbank model into the birth–death-sampling (BDS) [79–81] framework would offer a complementary approach to our coalescent framework. This approach would involve incorporating a multi-type BDS model [82–85] with a state-specific mutation rate model, similar to those in recent studies [86, 87], but tailored for the seedbank model.

## 6 Materials and Methods

### 6.1 Sampling from seedbank genealogical prior

We computed the empirical means and variances of *T*_MRCA_, *L*^(*a*)^, and *L*^(*d*)^ from MCMC samples targeting the seedbank prior. For each parameter and operator combination, we ran a chain for 100 million iterations, sampled every 10,000 iterations, and thus obtained 10,000 total samples. After discarding a 10% burn-in, we retained 9,000 posterior samples, from which we derived the empirical first and second moments of *T*_MRCA_, *L*^(*a*)^, and *L*^(*d*)^. The analytical recursive expressions for their expectations and variances, as functions of seedbank model parameters (*θ, c*, and *K*), were obtained from Blath et al. [9] and evaluated numerically.

### 6.2 Synthetic data generation

Synthetic molecular data were generated using our custom scripts in two stages: (1) simulating genealogies and (2) conditional on the simulated genealogies, sampling molecular sequences by superimposing a mutation process on the genealogy.

First, we simulated seedbank genealogies under the seedbank coalescent given specified seedbank model parameters and sample configurations. Our seedbank genealogy generation algorithm supports both isochronous and serial sampling schemes. Isochronous sampling specifies the initial number of active and dormant samples at *t* = 0. Serial sampling defines time intervals, each associated with a specified number of active and dormant samples within that interval. The waiting time until the next event was sampled from an exponential distribution with a rate equal to the sum of the coalescent and type-change rates, based on the current configuration of active and dormant lineages. Conditioned on this waiting time, an event type—coalescence, active-to-dormant transition, or dormant-to-active transition—was chosen with probability proportional to its respective rate. The affected lineage states and overall configuration were then updated accordingly. In the case of serial sampling, if a predefined sampling time occurred during the waiting interval, the sampling event took precedence over a coalescent or type-change event. This iterative process continued until only the genealogy coalesced into a single lineage representing the most recent common ancestor.

Next, given the simulated seedbank genealogy and the specified mutation model parameters, we generated molecular sequences using a modified implementation of the `TreeTime` package [88]. Starting from an ancestral root sequence with equal nucleotide frequencies, we evolved sequences along each branch in a preorder traversal under the HKY substitution model [65], with branch-specific mutation rates to reflect the active or dormant state of the lineage. The transition-to-transversion ratio (*κ*) was set to 3.0 across all simulations.

In our simulations, we considered both isochronous and serial sampling scenarios. Each dataset comprised 25 sequences, each 20,000 nucleotides in length. Under isochronous sampling, all 25 samples were active at *t* = 0. Under serial sampling, 20 active and 5 dormant samples were distributed across three time intervals, [0, 2), [2, 4), and [4, 6), with active samples divided into groups of (8, 8, 4) and dormant samples into groups of (2, 2, 1). All samples were placed randomly in their respective intervals. Under each of these four sampling configurations, six seedbank parameter sets were tested by varying the seedbank size *K* ∈ {1.0, 10.0} and the dormancy parameter *α* ∈ [0.1, 0.5, 0.99], while fixing *c* = 1.0, *θ* = 1.0, and *μ* = 0.01.

### 6.3 Inference from synthetic molecular data

We assigned LogNormal(0, 0.5), [0.375, 2.66] priors to both *c* and *θ*. The bracketed interval indicates the 95% interval for each prior. For *K*, we used LogNormal(0, 0.5), [0.375, 2.66] when *K* = 1.0 and LogNormal(2.5, 0.5), [4.57, 32.5] when *K* = 10.0. For *α*, we imposed Beta priors: Beta(1, 9), [0.00281, 0.336] for *α* = 0.1, Beta(10, 10), [0.289, 0.711] for *α* = 0.5, and Beta(9, 1), [0.664, 0.997] for *α* = 0.99. We assigned a LogNormal(−2.0, 2.0), [0.00269, 6.82] prior to *μ* and a LogNormal(1.0, 0.5), [1.02, 7.24] prior to *κ*. We ran MCMC chains for 100 million steps, sampling every 5,000 steps, with a 10% burn-in, yielding 18,000 posterior samples. For runs with slower convergence, joint tree inference with serial sampling, we increased the burn-in. Convergence and effective sample sizes (ESS) were assessed in `Tracer` v1.7.1, applying an ESS threshold of 200, except for tree height related parameters, where 100 was used.

For the joint inference of model parameters and the seedbank genealogy, we initialized a random seedbank genealogy from the leaf set using `SeedbankTreeInitializer`. We employed the seedbank tree operators (Sections 3.4.2 and S3)—tree-height scaling, Wilson–Balding, narrow and wide subtree exchange, node-height shift, and edge-recolor operators—to traverse the state space of seedbank genealogies. For inference under a fixed known genealogy, the true reduced genealogy was provided in Newick format into the `SeedbankTreeFromNewick` class. In both the joint and the fixed genealogy inferences, the edge-recolor operator (Section S3.6) was used to propose new configurations of lineage states (active or dormant) along edges while preserving constraints imposed by the seedbank coalescent model.

For inference under the misspecified model, we estimated {*T*_MRCA_, *μ*_*a*_, *κ*} from synthetic data previously generated under the seedbank coalescent, using the Kingman coalescent with a constant effective population size, the HKY substitution model, and a strict molecular clock. We placed an inverse prior on *θ*, a uniform prior on *μ*, and a LogNormal(1, 1.25), [0.235, 31.5] prior on *κ*. We performed MCMC sampling for 10 million iterations, thinning every 1000 steps and discarding a 10% burn-in, yielding 9000 posterior samples per run.

### 6.4 Mtb genetic data analyses

The dataset of Colangeli et al. [43] consists of WGS of *Mtb* from ten individuals with known or putative epidemiological links, sampled over a multi-decade outbreak of the Rangipo strain in New Zealand. These samples span 20 years (1992–2011) and include three cases hypothesized to have undergone prolonged latency. Of the ten samples, two (sample IDs: *C*1 and *C*2) were derived from the same individual, and two (sample IDs: *A* and *T*) were from the same family. We retained only one sample with the earlier sampling date from each pair, resulting in eight independent samples.

We mapped the SNP data from Colangeli et al. [43] to the H37Rv reference genome (GCF_000195955.2; [89]) to obtain the WGS of the eight samples. Because the reported SNPs were annotated only at the gene level without specifying intragenic positions, each SNP was randomly assigned to a nucleotide within the corresponding gene that matched the reference allele, using positional information from the TB Genome Annotation Portal [90]. This random assignment does not affect our downstream inference, as our analysis assumes no linkage disequilibrium among loci. Sampling times were estimated from the reported year of active *Mtb* onset for each case, assuming the samples were collected in the same year as disease development.

### 6.5 Inference of *Mtb* pathogen dynamics

We assigned LogNormal priors to the parameters *c, K, θ, κ* and *μ*_*a*_, and a beta prior to *α*. The 95% interval for each prior distribution is denoted in brackets:

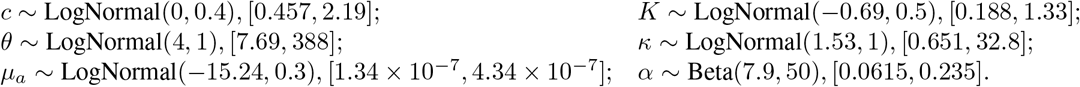

We ran five independent MCMC chains of 100 million steps each, discarding a 10% burn-in. We assessed convergence in `Tracer` v1.7.1, with an ESS threshold of 1,000. The chains were combined using `LogCombiner` v2.7.6 and thinned to 8,480 samples. We summarized the posterior tree distribution using `TreeAnnotator` with the common ancestor heights option [91]. The MCC tree was visualized in `R` using the `ggtree` package [92, 93], coloring each edge by its median latent-state proportion λ.

## Data, Materials, and Software Availability

Our open-source software as a BEAST2 package is available on GitHub at BEAST-seedbank/SeedbankTree. The code used for analyzing both simulated and real data presented in this manuscript is available on Zenodo at 10.5281/zen-odo.14621998.

## Acknowledgements

LC was supported by the Ramon y Cajal 2022 RYC2022-038467-I, financed by MCIN/AEI/10.13039/501100011033 and FSE+. WTL, PX, and JK were supported by the National Institute of General Medical Sciences of the National Institutes of Health under award number R35GM156957. AGC and MTW were supported by the National Institute of Allergy and Infectious Diseases of the National Institutes of Health under award number P01AI159402.

