## Supplementary Information for "Bayesian phylodynamic inference of population dynamics with dormancy"

### Supplemental Information: Bayesian phylodynamic inference of population dynamics with dormancy

### SI Text

#### S1 Proof of Lemma 1

We introduce auxiliary nodes to prove Lemma 1. A description of these auxiliary nodes is given in the caption of Figure S1. The location of these auxiliary nodes is determined by the corresponding coloring $\psi_e$ assigned to each edge in the seedbank genealogy $\mathbf{g}$.

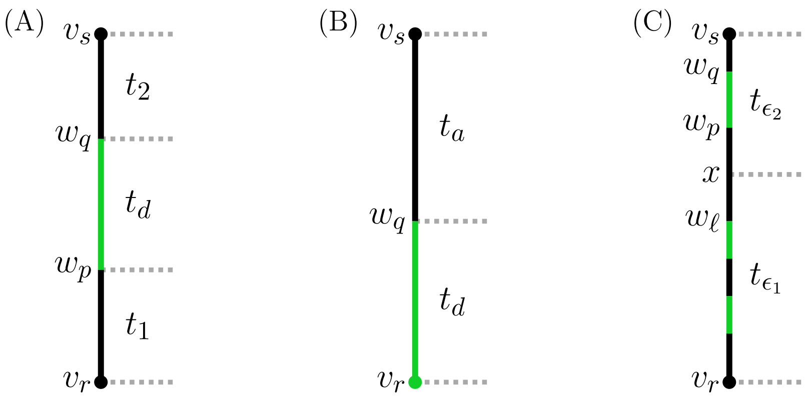


**Figure S1. Example of an edge $e = \{v_r, v_s\}$ with one or more dormant segments.** The black and green lines represent the active and dormant states, respectively. (A) An edge with a single dormant segment defined by two type-change events described by $\psi_e$. Here, $v_r, v_s \in S \cup C$, and let $w_p, w_q$ be the auxiliary type-change nodes corresponding to transitions from active to dormant ($a \to d$) and dormant to active ($d \to a$), respectively. The total length of the active state is $t_a = t_1 + t_2 = (1 - \lambda)t_e$, and the total length of the dormant state is $t_d = \lambda t_e$. (B) An edge with a single dormant segment defined by a single type-change event (again, captured by the corresponding $\psi_e$). In this case, edge $e$ is a terminal branch with $v_r \in S$ being a sample from the dormant population, and the auxiliary single type-change node $w_q$ represents the $d \to a$ transition. (C) An edge with multiple dormant segments. When edge $e$ contains $n + 1$ dormant segments, $w_\ell$ denotes the $d \to a$ auxiliary type-change node terminating the $n$-th dormant segment, and $w_p$ represents the $a \to d$ auxiliary type-change node initiating the $n + 1$-th dormant segment. A further auxiliary node $x \notin S \cup C$, placed at the midpoint between $w_\ell$ and $w_p$, partitions edge $e$ into two sub-edges: $\epsilon_1 = \langle v_r, x\rangle$ with length $t_{\epsilon_1}$ containing $n$ dormant segments, and $\epsilon_2 = \langle x, v_s\rangle$ with length $t_{\epsilon_2}$ containing one dormant segment.

**Lemma S1** (Transition probability matrix for an edge with a single dormant segment)**.** *Let $\mathbf{\Pi_a}$ and $\mathbf{\Pi_d}$ be the time-reversible substitution rate matrices for the active and dormant populations, respectively, with $\mathbf{\Pi_d} = \alpha\mathbf{\Pi_a}$ and $\alpha \in [0, 1]$, as defined in Section 3.2. Consider an edge $e = \langle v_r, v_s\rangle \in E$ of length $t_e$, where $v_r, v_s \in S \cup C$, which contains a single dormant segment occupying a fraction $\lambda$ of its length. The transition probability matrix $\mathbf{P_e}(t_e)$ for edge $e$ is given by:*

$$\mathbf{P_e}(t_e) = e^{\mathbf{\Pi_a}[1-(1-\alpha)\lambda]t_e}. \tag{S1}$$

*Proof.* We first consider the case where the single dormant segment is defined by two auxiliary type-change events at nodes $w_p, w_q$, as shown in Figure S1(A). The transition probability matrix along the edge $e$ is computed as:

$$\mathbf{P_e}(t_e) = e^{\mathbf{\Pi_a}t_2}e^{\mathbf{\Pi_d}t_d}e^{\mathbf{\Pi_a}t_1} = e^{\mathbf{\Pi_a}(t_1+t_2+\alpha t_d)} = e^{\mathbf{\Pi_a}(t_a+\alpha t_d)} = e^{\mathbf{\Pi_a}[1-(1-\alpha)\lambda]t_e},$$

where $t_a = t_1 + t_2 = (1 - \lambda)t_e$, and $t_d = \lambda t_e$ by definition. The second equality follows from $\mathbf{\Pi_d} = \alpha\mathbf{\Pi_a}$ and the property $e^{\mathbf{X}}e^{\mathbf{Y}} = e^{\mathbf{X}+\mathbf{Y}}$ for any commuting matrices $\mathbf{X}$ and $\mathbf{Y}$.

When the single dormant segment is defined by a single type-change event at auxiliary node $w_q$, with edge $e$ as a terminal branch containing a dormant sample $v_r \in S$, as shown in Figure S1(B), the transition probability matrix along the edge $e$ is computed as:

$$\mathbf{P_e}(t_e) = e^{\mathbf{\Pi_a}t_a}e^{\mathbf{\Pi_d}t_d} = e^{\mathbf{\Pi_a}(t_a+\alpha t_d)} = e^{\mathbf{\Pi_a}[1-(1-\alpha)\lambda]t_e}.$$

Thus, the overall transition probability matrix for edge $e$ containing a single dormant segment is $\mathbf{P_e}(t_e) = e^{\mathbf{\Pi_a}[1-(1-\alpha)\lambda]t_e}$.

□

**Lemma S2** (Transition probability matrix for an edge with multiple dormant segments)**.** *Consider an edge $e = \langle v_r, v_s\rangle \in E$ of length $t_e$, where $v_r, v_s \in S \cup C$, containing multiple dormant segments that occupy a total fraction $\lambda$ of its length. The transition probability matrix $\mathbf{P_e}(t_e)$ for edge $e$ is given by:*

$$\mathbf{P_e}(t_e) = e^{\mathbf{\Pi}_{\boldsymbol{a}}[1-(1-\alpha)\lambda]t_e}. \tag{S2}$$

*Proof.* We proceed by induction on the number of dormant segments $n$ in edge $e$. In the base case of a single dormant segment ($n = 1$), the lemma holds as established in Lemma S1. Assuming that the lemma holds for an edge with $n \geq 1$ dormant segments, we consider an edge $e = \langle v_r, v_s\rangle$ with $n + 1$ dormant segments and prove that the lemma remains valid in this case.

Let $w_\ell$ denote the auxiliary node corresponding to the $d \to a$ type-change at the end of the $n$-th dormant segment, and let $w_p, w_q$ be the auxiliary type-change nodes defining the $(n + 1)$-th dormant segment; see Figure S1(C). We partition edge $e$ into two sub-edges by introducing a further auxiliary node $x \notin S \cup C$ at the midpoint of the active portion between $w_\ell$ and $w_p$, as shown in Figure S1(C). Let $\epsilon_1 = \langle v_r, x\rangle$ and $\epsilon_2 = \langle x, v_s\rangle$, with respective lengths $t_{\epsilon_1}$ and $t_{\epsilon_2}$. Then, $\epsilon_1$ contains $n$ dormant segments, and $\epsilon_2$ contains one dormant segment. Define $\lambda_1$ and $\lambda_2$ as the fractions of the total dormant lengths within $\epsilon_1$ and $\epsilon_2$, respectively.

The transition probability matrix for edge $e$ with $n + 1$ dormant segment is:

$$\begin{aligned}\mathbf{P_e}(t_e) &= \mathbf{P}_{\boldsymbol{\epsilon_2}}(t_{\epsilon_2})\mathbf{P}_{\boldsymbol{\epsilon_1}}(t_{\epsilon_1}) \\ &= e^{\mathbf{\Pi}_{\boldsymbol{a}}[1-(1-\alpha)\lambda_2]t_{\epsilon_2}} e^{\mathbf{\Pi}_{\boldsymbol{a}}[1-(1-\alpha)\lambda_1]t_{\epsilon_1}} \\ &= e^{\mathbf{\Pi}_{\boldsymbol{a}}\left[(t_{\epsilon_1}+t_{\epsilon_2})-(1-\alpha)(\lambda_1 t_{\epsilon_1}+\lambda_2 t_{\epsilon_2})\right]} \\ &= e^{\mathbf{\Pi}_{\boldsymbol{a}}[1-(1-\alpha)\lambda]t_e},\end{aligned}$$

where the second equality follows from Lemma S1 and the induction hypothesis for $n$. The third equality uses $t_e = t_{\epsilon_1} + t_{\epsilon_2}$ and $\lambda = \dfrac{\lambda_1 t_{\epsilon_1} + \lambda_2 t_{\epsilon_2}}{t_e}$, the total fraction of dormant length within edge $e$. This completes the proof.

□

By Lemma S2, the effective substitution rate matrix for any edge $e_i \in E$, $i \in \{1, \ldots, 2n - 2\}$, of the seedbank genealogy can be defined as:

$$\mathbf{\Pi}_{\boldsymbol{i}}^{\text{eff}} = [1 - (1 - \alpha)\lambda_i]\, \mathbf{\Pi}_{\boldsymbol{a}}.$$

#### S2 Proof of Theorem 2

**Overview of Felsenstein's pruning algorithm**: We consider a time-reversible continuous-time Markov chain (CTMC) model for molecular evolution [1, 2], characterized by the substitution rate matrix $\mathbf{\Pi}$. Let $\Sigma$ be the state space for sequence data, represented as a totally ordered set under lexicographic order; for example, for nucleotide sequences, $\Sigma = \{A, C, G, T\}$ with the total order $A < C < G < T$. We denote the $j$-th element of $\Sigma$ as $\sigma_j$. As defined in Section 3.1, $S$ represents the $n$ leaf nodes from the sampling events, and $C$ denotes the $n - 1$ internal (ancestral) nodes from the coalescent events between active lineages, including the root node. Let $L^{(i)}_{v_k}(\sigma_j)$ denote the conditional likelihood of the subtree rooted at node $v_k \in S \cup C$, given that $v_k$ is in state $\sigma_j \in \Sigma$ at site $i$, where $k \in [|S \cup C|]$ and $j \in [|\Sigma|]$.

Felsenstein's pruning algorithm [3] is a dynamic programming method for computing $L^{(i)}_{v_k}(\sigma_j)$ by traversing tree nodes in a specific order, such as post-order traversal. Let $Y^{(i)}_{v_k}$ represent the state at site $i$ of the observed sequence data at leaf node $v_k$. The initialization condition for each leaf node $v_k \in S$, with $k = 1, \ldots, n$, is given by:

$$L^{(i)}_{v_k}(\sigma_j) = \begin{cases} 1 & \text{if} \quad \sigma_j = Y^{(i)}_{v_k} \\ 0 & \text{if} \quad \sigma_j \neq Y^{(i)}_{v_k} \end{cases}. \tag{S3}$$

At each internal node $v_k \in C$ for $k = n+1, \ldots, 2n-1$, the conditional likelihood $L_{v_k}^{(i)}(\sigma_q)$ is computed recursively using its left ($v_\ell$) and right ($v_r$) direct child nodes:

$$L_{v_k}^{(i)}(\sigma_q) = \left( \sum_{j=1}^{|\Sigma|} P(\sigma_j \mid \sigma_q, t_\ell) L_{v_\ell}^{(i)}(\sigma_j) \right) \times \left( \sum_{j=1}^{|\Sigma|} P(\sigma_j \mid \sigma_q, t_r) L_{v_r}^{(i)}(\sigma_j) \right), \tag{S4}$$

where $t_\ell$ and $t_r$ are the branch lengths from node $v_k$ to its child nodes $v_\ell$ and $v_r$, respectively. $P(\sigma_j \mid \sigma_q, t)$ denotes the transition probability from state $\sigma_q$ to state $\sigma_j$ over time $t$, computed under the substitution model $\mathbf{\Pi}$. The algorithm terminates at the root, computing the overall likelihood of the tree for the sequence data $\mathbf{Y}^{(i)}$ at site $i$:

$$P(\mathbf{Y}^{(i)} \mid \mathbf{g}, \mathbf{\Pi}) = \sum_{j=1}^{|\Sigma|} f_{\sigma_j} L_{\text{root}}^{(i)}(\sigma_j), \tag{S5}$$

where $f_{\sigma_j}$ denotes the stationary frequency of state $\sigma_j$.

**Proof of Theorem 2:**

*Proof.* We prove the theorem by induction on the nodes $v_k \in S \cup C$. The base case for leaf nodes $v_k \in S$ follows from Eq. S3. Assuming the theorem holds for all descendants of an internal node $v_k \in C$, we proceed to prove it for $v_k$. Let $v_\ell$ and $v_r$ be the left and right child nodes of $v_k$, respectively, with corresponding edges $e_\ell = \{v_k, v_\ell\} \in E$ and $e_r = \{v_k, v_r\} \in E$, and edge lengths $t_\ell$ and $t_r$. Denote $\mathbf{L}_{v_k}^{(i)}$ as a column vector of length $|\Sigma|$, where its $j$-th element is $L_{v_k}^{(i)}(\sigma_j)$. We express the transition probability matrices as left stochastic matrices. The likelihood at node $v_k$ is then computed recursively using Eq. S4:

$$\mathbf{L}_{v_k}^{(i)} = \left( \mathbf{P}_{\mathbf{e}_\ell}(t_\ell) \mathbf{L}_{v_\ell}^{(i)} \right) \odot \left( \mathbf{P}_{\mathbf{e_r}}(t_r) \mathbf{L}_{v_r}^{(i)} \right),$$

where $\odot$ represents the Hadamard product. By Lemma S2 and the induction hypothesis, $\mathbf{L}_{v_k}^{(i)}$ can be computed using the branch-specific effective rate matrices. By induction through post-order traversal of the seedbank genealogy $\mathbf{g}$, this dependence extends to the conditional likelihoods at all nodes of $\mathbf{g}$. Thus, the overall likelihood of the seedbank genealogy $P(\mathbf{Y} \mid \mathbf{g}, \mathbf{\Pi}_{\boldsymbol{a}}, \mathbf{\Pi}_{\boldsymbol{d}})$, as given by Eq. S5, is equal to the phylogenetic likelihood of $\tilde{\mathbf{g}}$ with branch-specific effective substitution rate matrices as defined in Lemma 1. It therefore depends only on the substitution model parameters $\{\alpha, \mu_a, \mathbf{Q}\}$, edge lengths $\boldsymbol{\ell}$, and dormant state proportions $\boldsymbol{\lambda}$, confirming the main theorem. □

#### S3 MCMC transition kernels

We provide a description of the Metropolis-Hastings proposals used within the MCMC framework, along with their corresponding acceptance probabilities. The description includes both established moves from the literature, as well as modifications necessary for accommodating the seedbank coalescent.

The scheme followed is the reversible jump MCMC [4], which is standard in the structured coalescent literature (e.g., [5, 6]). Suppose that at iteration $t$ the chain is at a generic state $s$. The next state is determined as follows. A move (operator) $k$ is chosen with probability $r(k)$ from a discrete set of operators. Then, a random number $u$ is sampled from a density $g$. The proposed state of the chain $s^*$ is constructed via a suitable deterministic invertible map $h$ such that $(s^*, u^*) = h(s, u)$ where $u^*$ are random numbers generated by an operator $k^*$ from a known density $g^*$, required to reverse the move from $(s^*, u^*)$ to $(s, u)$. Following Green [4], the move is accepted with probability:

$$\alpha(s, s^*) = \min \left\{ 1, \frac{\pi(s^*)}{\pi(s)} \frac{r(k^*) g^*(u^*)}{r(k) g(u)} \left| \frac{\partial(s^*, u^*)}{\partial(s, u)} \right| \right\},$$

where $\pi$ denotes the target distribution, and the last term represents the Jacobian of the transformation $h$. While the acceptance ratio is generally used for a trans-dimensional move, it is valid in general (see Green [4] for more details) and is fully equivalent to a standard Metropolis–Hastings ratio when there is no trans-dimensional move.

In what follows, we describe the one-step-ahead proposal of our operator. For simplicity, we omit the time index on the variables. In line with standard practice in coalescent-based inference, we refer to the following quantity as the Hastings-Green ratio:

$$HGR(s, s^*) = \frac{r(k^*)g^*(u^*)}{r(k)g(u)} \left| \frac{\partial(s^*, u^*)}{\partial(s, u)} \right|,$$

where the numerator is the transition probability from $(s^*, u^*)$ to $(s, u)$, and the denominator corresponds to the reversed move.

##### S3.1 Parameter scaler [`ScaleOperator`]

This is the widely used parameter operator in base `BEAST2`, simply called `ScaleOperator`. We apply it to $c$, $K$, $\theta$, $\mu_a$, $\alpha$, and $\mathbf{Q}$. For a generic state $s$, draw $x \sim \mathcal{U}(f, 1/f)$, with $f \in (0, 1)$, and propose $s^* = s\,x$. The HGR is $1/x$. The proposal corresponds to a random walk on a log transformation of the parameter.

##### S3.2 Seedbank tree scaler [`SeedbankTreeScale`]

`SeedbankTreeScale` corresponds to the `MultiTypeTreeScaler` of [5, 6]. The full genealogy is rescaled, which involves rescaling the node heights of all coalescent and type-transition nodes. The scaling factor is the same for the whole genealogy.

Draw a multiplier $x$ uniformly at random from $[f, \frac{1}{f})$. Then, iterate through all the internal coalescent and transition nodes and scale the corresponding height. The proposal may be invalid (child older than parent), in which case, the move is rejected. Otherwise, the HGR is $\frac{x^{|C|+|M|}}{x^2}$.

##### S3.3 Node-shift retype [`NodeShiftRetype`]

`NodeShiftRetype` is one of the two-step operators proposed by [6]. Let $u_p$, $u_l$, and $u_r$ be the parent, left child, and right child of a node $u$ respectively.

First, select random coalescent node $i \in C$. If $i$ is the root, the new root height is found by rescaling the difference between $t(i)$ and $\max(t(i_l), t(i_r))$. Then propose new state $\psi_e^* \in M$ for the edges $\langle i_l, i \rangle$ and $\langle i_r, i \rangle$ by sampling from the endpoint conditioned CTMC $\psi_e(t)$. If $i$ is not the root, disconnect $i$ from $i_p$, $i_l$, and $i_r$, and then uniformly draw a new location for $i$ from $[\max(t(i_l), t(i_r)), t(i_p)]$. Then, propose new $\psi_e^* \in M$ for the edges $\langle i_l, i \rangle$, $\langle i_r, i \rangle$, and $\langle i, i_p \rangle$. In the last step, update $\boldsymbol{\lambda}^*$ and $\boldsymbol{\rho}^*$ by updating all edges involved in the proposal.

Let $E^* \subset E$ denote the subset of the edges involved in the proposal, and let $s$ represent all variables involved in the proposal. The HGR is

$$HGR(s, s^*) = \begin{cases} \dfrac{\prod_{e \in E^*} P(\psi_e)}{\prod_{e \in E^*} P(\psi_e^*)} \dfrac{1}{x} & \text{if } i \text{ is the root} \\[2ex] \dfrac{\prod_{e \in E^*} P(\psi_e)}{\prod_{e \in E^*} P(\psi_e^*)} & \text{otherwise} \end{cases}, \tag{S6}$$

where $P(\psi_e)$ and $P(\psi_e^*)$ are computed following the description of the uniformization-based scheme in Section 3.4.1.

##### S3.4 Modified subtree exchange with coloring [`TypedSubtreeExchange`]

`TypedSubtreeExchange` is one of the two-step operators proposed by [6]. Let $u_p$ and $u_s$ be the parent and sibling of a node $u$, respectively. Randomly select samples or coalescent nodes $i$ and $j$, where $j$ is not $Sib(i, \tilde{\mathbf{g}})$. If $t(i) > t(j_p)$ or $t(j) > t(i_p)$, reject the proposal. Swap $i$ and $j$, and then propose a new state $\psi_e^* \in M$ for the edges $\langle i, i_p \rangle$ and $\langle j, j_p \rangle$. Update $\boldsymbol{\lambda}^*$ and $\boldsymbol{\rho}^*$ by updating all edges involved in the proposal. The move is accepted with probability:

$$HGR(s, s^*) = \frac{\prod_{e \in E^*} P(\psi_e)}{\prod_{e \in E^*} P(\psi_e^*)}.$$

##### S3.5 Modified Wilson-Balding with coloring [`TypedWilsonBalding`]

`TypedWilsonBalding` is an operator proposed by [6]. It can only be used on trees with more than three leaf nodes. Let $u_p$, $u_{gp}$, and $u_s$ be the parent, grandparent, and sibling of a node $u$, respectively. Randomly select a sample or coalescent node $i$ such that $i$ is not the root and $i_p$ is not the root. Randomly pick another sample or coalescent node, $j$, such that $t(j_p) > t(j)$. Additionally, $j$ is not the root, $j_p$ is not the root, and $j$ is not $i_p$ or $i_s$.

Detach $i$ from the tree, replacing $\langle i_s, i_p \rangle$ and $\langle i_p, i_{gp} \rangle$ with $\langle i_s, i_{gp} \rangle$. Now select a new location uniformly at random from $[\max(t(i), t(j)), t(j_p)]$ to reattach $i$, replacing $\langle j, j_p \rangle$ with $\langle j, i_p \rangle$ and $\langle i_p, j_p \rangle$. If the type of $\langle j, j_p \rangle$ at the attachment point is dormant, then reject the proposal. Finally, propose a new state $\psi_e^* \in M$ for the edge $\langle i, i_p \rangle$, and update $\boldsymbol{\lambda}^*$ and $\boldsymbol{\rho}^*$ for all edges involved in the proposal. The move is accepted with probability:

$$HGR(s, s^*) = \frac{\prod_{e \in E^*} P(\psi_e)}{\prod_{e \in E^*} P(\psi_e^*)} \times \frac{t(j_p) - max(t(i), t(j))}{t(i_{gp}) - max(t(i), t(i_s))}.$$

##### S3.6 Edge type change [`EdgeRecolor`]

`EdgeRecolor` is an operator specific to the seedbank coalescent. It selects an edge at random and samples a new coloring using the uniformization-based scheme described in Section 3.4.1. The move is accepted with probability:

$$HGR(s, s^*) = \frac{P(\psi_e)}{P(\psi_e^*)}.$$

#### SI Tables

**Table S1. Summary of notations and definitions.**

| Symbol | Definition |
|---|---|
| $a$ | Active type |
| $d$ | Dormant type |
| $\mathcal{P}_k^{\{a,d\}}$ | Space of typed partition of $[k]$, where each block is labeled as either $a$ or $d$ |
| $N_a$ | Effective population size of active population |
| $N_d$ | Effective population size of dormant population |
| $c$ | Transition rate from active to dormant |
| $K$ | Relative size of active population compared to dormant population such that $N_a = KN_d$ |
| $S$ | Set of leaf nodes from sampling events |
| $C$ | Set of coalescent nodes |
| $n$ | Number of samples |
| $n_c$ | Number of coalescent events |
| $V$ | Set of coalescent and sampling nodes |
| $E$ | Set of edges defined by nodes in $V$ |
| $\psi_{e_i}$ | Transitions ($a \to d$ or $d \to a$) occurring on edge $e_i \in E$, along with their event times |
| $M$ | Set of sample paths of type-change processes: $\{\psi_{e_1}, \ldots, \psi_{e_{2n-2}}\}$ |
| $\mathbf{u}$ | Vector of ordered event times measured in calendar time units |
| $\mathbf{g}$ | Seedbank genealogy: $\mathbf{g} = (V, E, \mathbf{u}, M)$ |
| $\tilde{\mathbf{g}}$ | Reduced genealogy: $\tilde{\mathbf{g}} = (V, E, \mathbf{u})$ |
| $n_{da}$ | Number of dormant-to-active type-changes in $\mathbf{g}$ |
| $n_{ad}$ | Number of active-to-dormant type-changes in $\mathbf{g}$ |
| $\theta$ | Scaling factor converting coalescent time units to calendar time, $\theta = N_a \tau_g$ |
| $\tau_g$ | Generation time in calendar units |
| $\Delta_i$ | Length of time interval $(u_i, u_{i+1})$ |
| $k_{i,a}$ | Number of active lineages during time interval $(u_i, u_{i+1})$ |
| $k_{i,d}$ | Number of dormant lineages during time interval $(u_i, u_{i+1})$ |
| $\mu_a$ | Mutation rate per site per unit calendar time of active population |
| $\mu_d$ | Mutation rate per site per unit calendar time of dormant population |
| $\mathbf{Q}$ | Normalized base substitution rate matrix |
| $\mathbf{\Pi_a}$ | Substitution rate matrix for active population |
| $\mathbf{\Pi_d}$ | Substitution rate matrix for dormant population |
| $\alpha$ | Ratio of mutation rates between dormant and active populations |
| $\ell_i$ | Edge length of edge $e_i$ |
| $\lambda_i$ | Proportion of dormancy in edge $e_i$ |
| $\xi_i$ | Branch-specific scaling factor, $\xi_i := 1 - (1 - \alpha)\lambda_i$ |
| $\rho_i$ | Branch-rate multiplier, $\rho_i := \xi_i \mu_a$ |
| $\mathbf{\Pi_i^{eff}}$ | Branch-specific effective substitution rate matrix of edge $e_i$ |
| $\mathbf{Y}$ | Aligned molecular sequences, along with their sampling times and states |
| $\kappa$ | transition/transversion bias |

**Table S2. Comparison of the analytic first and second moments of tree height between seedbank coalescent and two-population structured coalescent models.** (A) Analytic expectation of tree height, $\mathbb{E}[T_{\text{MRCA}}]$. (B) Analytic variance of tree height, Var$[T_{\text{MRCA}}]$. The analytic moments for the seedbank coalescent were computed by numerically evaluating the recursive formula in Blath et al. [7]. The effective population size of the active population ($N_a$) was fixed at 1, and the seedbank model parameters $c$ and $K$ were varied as specified in the table. The closed-form expressions for the structured coalescent are provided in Wakeley [8]. For fair comparisons between the two models, analogous population parameters were matched: in the two-population structured coalescent, the effective populations sizes were set to $N_1 = N_a$ and $N_2 = N_a/K$, corresponding to the active and dormant populations in the seedbank coalescent, respectively.

**(A) Analytic expectation of tree height, $\mathbb{E}[\boldsymbol{T}_{\textbf{MRCA}}]$.**

| | | $n=2$ | | $n=10$ | | $n=100$ | |
|---|---|---|---|---|---|---|---|
| $c$ | $K$ | Seedbank | Structured | Seedbank | Structured | Seedbank | Structured |
| 0.1 | 0.1 | 121 | 4.182 | 322.7 | 10.12 | 530.9 | 15.12 |
| 0.5 | 0.1 | 121 | 6.909 | 281.4 | 15.17 | 378.2 | 19.19 |
| 1.0 | 0.1 | 121 | 8.273 | 259.4 | 17.07 | 321.4 | 20.29 |
| 5.0 | 0.1 | 121 | 10.26 | 228.3 | 19.14 | 259.0 | 21.36 |
| 10.0 | 0.1 | 121 | 10.61 | 223.2 | 19.45 | 249.6 | 21.54 |
| 0.1 | 1.0 | 4 | 2.000 | 9.671 | 4.369 | 14.97 | 5.953 |
| 0.5 | 1.0 | 4 | 2.000 | 8.564 | 3.957 | 11.32 | 4.626 |
| 1.0 | 1.0 | 4 | 2.000 | 8.071 | 3.803 | 10.02 | 4.297 |
| 5.0 | 1.0 | 4 | 2.000 | 7.425 | 3.641 | 8.480 | 4.022 |
| 10.0 | 1.0 | 4 | 2.000 | 7.317 | 3.620 | 8.221 | 3.989 |
| 0.1 | 10.0 | 1.21 | 1.168 | 2.279 | 2.180 | 2.697 | 2.512 |
| 0.5 | 10.0 | 1.21 | 1.141 | 2.210 | 2.071 | 2.491 | 2.291 |
| 1.0 | 10.0 | 1.21 | 1.127 | 2.195 | 2.036 | 2.448 | 2.244 |
| 5.0 | 10.0 | 1.21 | 1.107 | 2.182 | 1.994 | 2.407 | 2.194 |
| 10.0 | 10.0 | 1.21 | 1.104 | 2.180 | 1.987 | 2.402 | 2.186 |

**(B) Analytic variance of tree height, Var$[\boldsymbol{T}_{\textbf{MRCA}}]$.**

| | | $n=2$ | | $n=10$ | | $n=100$ | |
|---|---|---|---|---|---|---|---|
| $c$ | $K$ | Seedbank | Structured | Seedbank | Structured | Seedbank | Structured |
| 0.1 | 0.1 | 168600 | 30.71 | 397700 | 54.28 | 565800 | 60.98 |
| 0.5 | 0.1 | 45440 | 5.538 | 73000 | 4.214 | 75590 | 1.550 |
| 1.0 | 0.1 | 30040 | 2.982 | 40920 | 1.192 | 40990 | 0.108 |
| 5.0 | 0.1 | 17720 | 1.580 | 20930 | 0.026 | 20970 | 0.001 |
| 10.0 | 0.1 | 16180 | 1.492 | 18900 | 0.007 | 18920 | 0.0001 |
| 0.1 | 1.0 | 116.0 | 14.00 | 264.5 | 24.10 | 380.4 | 27.47 |
| 0.5 | 1.0 | 36.00 | 6.000 | 55.82 | 5.900 | 59.91 | 3.301 |
| 1.0 | 1.0 | 26.00 | 5.000 | 34.95 | 3.765 | 35.72 | 0.735 |
| 5.0 | 1.0 | 18.00 | 4.200 | 21.36 | 0.910 | 21.42 | 0.001 |
| 10.0 | 1.0 | 17.00 | 4.100 | 19.91 | 0.458 | 19.95 | 0.0002 |
| 0.1 | 10.0 | 1.915 | 1.799 | 2.636 | 2.358 | 2.962 | 2.601 |
| 0.5 | 10.0 | 1.554 | 1.511 | 1.839 | 1.820 | 1.851 | 2.185 |
| 1.0 | 10.0 | 1.509 | 1.474 | 1.763 | 1.789 | 1.767 | 2.231 |
| 5.0 | 10.0 | 1.473 | 1.443 | 1.708 | 1.784 | 1.711 | 2.169 |
| 10.0 | 10.0 | 1.469 | 1.439 | 1.702 | 1.785 | 1.704 | 2.091 |

**Table S3. Comparison of analytic and MCMC-based estimates of the first and second moments of tree height.** (A) Expectation of tree height, $\mathbb{E}[T_{\text{MRCA}}]$. (B) Variance of tree height, Var$[T_{\text{MRCA}}]$. This table compares the analytically derived the first and second moments of the time to the most recent common ancestor ($T_{\text{MRCA}}$) and the corresponding estimates from MCMC sampling across various parameter configurations of the seedbank coalescent model. MCMC estimates are based on sampling from the prior distribution of seedbank genealogies (Eq. 2), while the analytical values are from numerical evaluation of recursive formulas from Blath et al. [7]. Three sample sizes, $n = 2, 10, 100$, were examined, with all samples originating from the active population, and the seedbank model parameters ($c$ and $K$) were varied systematically. For $n = 2$, a single node-shift retype (NSR) operator or a combination of seedbank tree scaler (TS) and edge-recolor (RC) operators was employed. For larger sample sizes, $n = 10$ and $n = 100$, a complete set of operators was used.

**(A) Expectation of tree height, $\mathbb{E}[T_{\text{MRCA}}]$.**

| | | $n = 2$ | | | $n = 10$ | | $n = 100$ | |
|---|---|---|---|---|---|---|---|---|
| $c$ | $K$ | Analytic | NSR | TS+RC | Analytic | MCMC | Analytic | MCMC |
| 0.1 | 0.1 | 121 | 116.1 | 127.7 | 322.7 | 316.5 | 530.9 | 530.9 |
| 0.5 | 0.1 | 121 | 120.2 | 118.4 | 281.4 | 279.3 | 378.2 | 383.0 |
| 1.0 | 0.1 | 121 | 117.8 | 123.3 | 259.4 | 258.7 | 321.4 | 324.1 |
| 5.0 | 0.1 | 121 | 123.8 | 120.2 | 228.3 | 230.2 | 259.0 | 260.8 |
| 10.0 | 0.1 | 121 | 120.5 | 112.7 | 223.2 | 223.8 | 249.6 | 248.3 |
| 0.1 | 1.0 | 4 | 4.172 | 3.867 | 9.671 | 9.784 | 14.97 | 16.12 |
| 0.5 | 1.0 | 4 | 3.957 | 4.025 | 8.564 | 8.612 | 11.32 | 11.32 |
| 1.0 | 1.0 | 4 | 4.072 | 4.023 | 8.071 | 8.090 | 10.02 | 10.03 |
| 5.0 | 1.0 | 4 | 4.015 | 4.018 | 7.425 | 7.407 | 8.480 | 8.701 |
| 10.0 | 1.0 | 4 | 3.991 | 3.994 | 7.317 | 7.341 | 8.221 | 8.192 |
| 0.1 | 10.0 | 1.21 | 1.218 | 1.205 | 2.279 | 2.282 | 2.697 | 2.746 |
| 0.5 | 10.0 | 1.21 | 1.211 | 1.227 | 2.210 | 2.206 | 2.491 | 2.502 |
| 1.0 | 10.0 | 1.21 | 1.212 | 1.202 | 2.195 | 2.187 | 2.448 | 2.454 |
| 5.0 | 10.0 | 1.21 | 1.211 | 1.209 | 2.182 | 2.168 | 2.407 | 2.400 |
| 10.0 | 10.0 | 1.21 | 1.206 | 1.205 | 2.180 | 2.153 | 2.402 | 2.406 |

**(B) Variance of tree height, Var$[T_{\text{MRCA}}]$.**

| | | $n = 2$ | | | $n = 10$ | | $n = 100$ | |
|---|---|---|---|---|---|---|---|---|
| $c$ | $K$ | Analytic | NSR | TS+RC | Analytic | MCMC | Analytic | MCMC |
| 0.1 | 0.1 | 168600 | 165800 | 173900 | 397700 | 375000 | 565800 | 603100 |
| 0.5 | 0.1 | 45440 | 42890 | 45160 | 73000 | 75530 | 75590 | 68450 |
| 1.0 | 0.1 | 30040 | 28820 | 30810 | 40920 | 41990 | 40990 | 46180 |
| 5.0 | 0.1 | 17720 | 18730 | 17040 | 20930 | 21050 | 20970 | 21710 |
| 10.0 | 0.1 | 16180 | 16190 | 16730 | 18900 | 19490 | 18920 | 18510 |
| 0.1 | 1.0 | 116.0 | 122.9 | 104.5 | 264.5 | 270.6 | 380.4 | 425.8 |
| 0.5 | 1.0 | 36.00 | 35.97 | 35.06 | 55.82 | 56.78 | 59.91 | 59.74 |
| 1.0 | 1.0 | 26.00 | 26.97 | 25.11 | 34.95 | 34.70 | 35.72 | 35.10 |
| 5.0 | 1.0 | 18.00 | 17.73 | 18.48 | 21.36 | 20.77 | 21.42 | 23.72 |
| 10.0 | 1.0 | 17.00 | 16.56 | 17.05 | 19.91 | 20.09 | 19.95 | 19.66 |
| 0.1 | 10.0 | 1.915 | 1.901 | 1.905 | 2.636 | 2.599 | 2.962 | 3.115 |
| 0.5 | 10.0 | 1.554 | 1.532 | 1.600 | 1.839 | 1.803 | 1.851 | 1.840 |
| 1.0 | 10.0 | 1.509 | 1.534 | 1.516 | 1.763 | 1.703 | 1.767 | 1.734 |
| 5.0 | 10.0 | 1.473 | 1.499 | 1.497 | 1.708 | 1.671 | 1.711 | 1.679 |
| 10.0 | 10.0 | 1.469 | 1.480 | 1.461 | 1.702 | 1.626 | 1.704 | 1.631 |

**Table S4. Comparison of analytic and MCMC-based estimates of the expected total lengths of active lineages ($\mathbb{E}[L^{(a)}]$) and dormant lineages ($\mathbb{E}[L^{(d)}]$).** Each cell contains two rows: the first row presents $\mathbb{E}[L^{(a)}]$, and the second row shows $\mathbb{E}[L^{(d)}]$. The table structure mirrors that of Table S3.

| $c$ | $K$ | $n = 2$ | | | $n = 10$ | | $n = 100$ | |
|---|---|---|---|---|---|---|---|---|
| | | Analytic | NSR | TS+RC | Analytic | MCMC | Analytic | MCMC |
| 0.1 | 0.1 | 22.00 | 21.43 | 23.06 | 61.15 | 60.49 | 105.3 | 112.2 |
| | | 220.0 | 210.8 | 232.3 | 611.5 | 600.2 | 1053 | 1132 |
| 0.5 | 0.1 | 22.00 | 21.84 | 21.50 | 59.27 | 58.87 | 92.93 | 94.63 |
| | | 220.0 | 218.5 | 215.4 | 592.7 | 587.7 | 929.3 | 950.2 |
| 1.0 | 0.1 | 22.00 | 21.37 | 22.34 | 58.81 | 58.51 | 90.37 | 90.53 |
| | | 220.0 | 214.1 | 224.3 | 588.1 | 585.5 | 903.7 | 905.2 |
| 5.0 | 0.1 | 22.00 | 22.55 | 21.87 | 60.08 | 60.43 | 95.54 | 95.69 |
| | | 220.0 | 225.0 | 218.4 | 600.8 | 604.2 | 955.4 | 956.5 |
| 10.0 | 0.1 | 22.00 | 21.92 | 22.31 | 60.88 | 61.01 | 99.92 | 99.54 |
| | | 220.0 | 219.1 | 223.1 | 608.8 | 609.8 | 999.2 | 995.8 |
| 0.1 | 1.0 | 4.000 | 4.087 | 3.942 | 11.23 | 11.31 | 19.98 | 20.80 |
| | | 4.000 | 4.257 | 3.793 | 11.23 | 11.32 | 19.98 | 21.40 |
| 0.5 | 1.0 | 4.000 | 3.974 | 4.010 | 11.14 | 11.15 | 19.38 | 19.32 |
| | | 4.000 | 3.94 | 4.039 | 11.14 | 11.21 | 19.37 | 19.22 |
| 1.0 | 1.0 | 4.000 | 4.081 | 4.014 | 11.15 | 11.18 | 19.40 | 19.48 |
| | | 4.000 | 4.064 | 4.032 | 11.15 | 11.17 | 19.40 | 19.48 |
| 5.0 | 1.0 | 4.000 | 4.027 | 4.022 | 11.24 | 11.20 | 19.92 | 20.11 |
| | | 4.000 | 4.003 | 4.014 | 11.24 | 11.19 | 19.92 | 20.14 |
| 10.0 | 1.0 | 4.000 | 3.997 | 3.999 | 11.27 | 11.33 | 20.16 | 20.10 |
| | | 4.000 | 3.985 | 3.989 | 11.27 | 11.34 | 20.16 | 20.13 |
| 0.1 | 10.0 | 2.200 | 2.221 | 2.194 | 6.221 | 6.234 | 11.36 | 11.45 |
| | | 0.2200 | 0.2145 | 0.2153 | 0.6221 | 0.6301 | 1.136 | 1.160 |
| 0.5 | 10.0 | 2.200 | 2.202 | 2.230 | 6.222 | 6.215 | 11.37 | 11.39 |
| | | 0.2200 | 0.2202 | 0.2234 | 0.6222 | 0.6165 | 1.137 | 1.145 |
| 1.0 | 10.0 | 2.200 | 2.201 | 2.188 | 6.222 | 6.210 | 11.38 | 11.41 |
| | | 0.2200 | 0.2224 | 0.2162 | 0.6222 | 0.6182 | 1.138 | 1.147 |
| 5.0 | 10.0 | 2.200 | 2.202 | 2.197 | 6.223 | 6.211 | 11.39 | 11.36 |
| | | 0.2200 | 0.2196 | 0.2202 | 0.6223 | 0.6198 | 1.139 | 1.131 |
| 10.0 | 10.0 | 2.200 | 2.193 | 2.190 | 6.224 | 6.174 | 11.39 | 11.40 |
| | | 0.2200 | 0.2192 | 0.2190 | 0.6224 | 0.6181 | 1.139 | 1.141 |

**Table S5. Comparison of analytic and MCMC-based estimates of the variance of total lengths of active lineages (Var$[L^{(a)}]$) and dormant lineages (Var$[L^{(d)}]$).** Each cell contains two rows: the first row presents Var$[L^{(a)}]$, and the second row shows Var$[L^{(d)}]$. The table structure mirrors that of Table S3.

| | | $n = 2$ | | | $n = 10$ | | $n = 100$ | |
|---|---|---|---|---|---|---|---|---|
| $c$ | $K$ | Analytic | NSR | TS+RC | Analytic | MCMC | Analytic | MCMC |
| 0.1 | 0.1 | 4484<br>572400 | 4518<br>561200 | 4584<br>590600 | 11080<br>1450000 | 10700<br>1368000 | 16490<br>2225000 | 17310<br>2378000 |
| 0.5 | 0.1 | 1284<br>153200 | 1212<br>144600 | 1255<br>152500 | 2424<br>308400 | 2493<br>319300 | 2746<br>374400 | 2569<br>351100 |
| 1.0 | 0.1 | 884.0<br>100800 | 851.4<br>96690 | 898.0<br>103500 | 1500<br>182900 | 1528<br>186700 | 1631<br>212100 | 1818<br>235100 |
| 5.0 | 0.1 | 564.0<br>58880 | 598.2<br>62220 | 544.1<br>56590 | 879.8<br>94760 | 876.0<br>94740 | 942.2<br>104900 | 957.0<br>107600 |
| 10.0 | 0.1 | 524.0<br>53640 | 524.1<br>53660 | 542.5<br>55440 | 811.8<br>84620 | 839.3<br>86980 | 866.7<br>92290 | 853.3<br>91320 |
| 0.1 | 1.0 | 56.00<br>216.0 | 58.46<br>230.6 | 52.32<br>192.3 | 128.7<br>574.0 | 131.1<br>572.4 | 187.1<br>957.3 | 210.7<br>1018 |
| 0.5 | 1.0 | 24.00<br>56.00 | 24.01<br>56.01 | 23.43<br>54.60 | 42.78<br>130.4 | 43.04<br>133.4 | 49.85<br>194.6 | 49.76<br>189.8 |
| 1.0 | 1.0 | 20.00<br>36.00 | 21.07<br>36.87 | 19.58<br>34.44 | 33.32<br>77.21 | 33.30<br>76.00 | 37.47<br>110.1 | 37.55<br>109.8 |
| 5.0 | 1.0 | 16.80<br>20.00 | 16.57<br>19.68 | 17.28<br>20.52 | 26.35<br>35.28 | 25.83<br>34.55 | 28.55<br>43.86 | 30.47<br>46.46 |
| 10.0 | 1.0 | 16.40<br>18.00 | 15.99<br>17.52 | 16.42<br>18.10 | 25.50<br>29.99 | 25.90<br>30.62 | 27.40<br>35.24 | 26.87<br>35.05 |
| 0.1 | 10.0 | 5.240<br>0.5724 | 5.297<br>0.5352 | 5.300<br>0.5455 | 8.538<br>1.554 | 8.406<br>1.580 | 9.748<br>2.765 | 10.16<br>2.829 |
| 0.5 | 10.0 | 4.920<br>0.1532 | 4.839<br>0.1541 | 5.068<br>0.1606 | 7.667<br>0.3705 | 7.607<br>0.3578 | 8.278<br>0.6170 | 8.272<br>0.6164 |
| 1.0 | 10.0 | 4.880<br>0.1008 | 4.939<br>0.1056 | 4.902<br>0.1022 | 7.560<br>0.2226 | 7.391<br>0.2167 | 8.098<br>0.3486 | 8.034<br>0.3591 |
| 5.0 | 10.0 | 4.848<br>0.0589 | 4.931<br>0.0605 | 4.929<br>0.0594 | 7.474<br>0.1042 | 7.348<br>0.1031 | 7.951<br>0.1332 | 7.969<br>0.1312 |
| 10.0 | 10.0 | 4.844<br>0.0536 | 4.874<br>0.0549 | 4.828<br>0.0523 | 7.463<br>0.0893 | 7.197<br>0.0871 | 7.932<br>0.1062 | 7.618<br>0.1026 |

**Table S6. Coverage, relative bias, and relative RMSE of the posterior estimates for key model parameters, based on 100 replicate simulations where only model parameters are inferred given the true reduced (uncolored) genealogy.** Table design follows Table 1 but excludes $T_{\text{MRCA}}$ because it is not applicable in this scenario.

| | Isochronous sampling | | | | | | Serial sampling | | | | | |
|---|---|---|---|---|---|---|---|---|---|---|---|---|
| | $K=1$ | | | $K=10$ | | | $K=1$ | | | $K=10$ | | |
| | $\alpha=0.1$ | $\alpha=0.5$ | $\alpha=0.99$ | $\alpha=0.1$ | $\alpha=0.5$ | $\alpha=0.99$ | $\alpha=0.1$ | $\alpha=0.5$ | $\alpha=0.99$ | $\alpha=0.1$ | $\alpha=0.5$ | $\alpha=0.99$ |
| | **Coverage** | | | | | | **Coverage** | | | | | |
| $c$ | 0.99 | 1.00 | 1.00 | 1.00 | 1.00 | 1.00 | 0.99 | 0.99 | 1.00 | 1.00 | 0.99 | 1.00 |
| $K$ | 0.99 | 1.00 | 0.99 | 1.00 | 1.00 | 1.00 | 0.97 | 0.98 | 1.00 | 1.00 | 1.00 | 1.00 |
| $\theta$ | 0.93 | 0.98 | 0.99 | 0.94 | 0.97 | 0.95 | 0.98 | 0.98 | 1.00 | 0.97 | 0.95 | 0.98 |
| $\kappa$ | 0.93 | 0.93 | 0.97 | 0.92 | 0.96 | 0.97 | 0.90 | 0.94 | 0.93 | 0.96 | 0.99 | 0.94 |
| $\alpha$ | | N/A | | | N/A | | 0.92 | 0.98 | 1.00 | 0.99 | 1.00 | 1.00 |
| $\mu_a$ | | N/A | | | N/A | | 0.92 | 0.97 | 0.93 | 0.94 | 0.94 | 0.94 |
| | **Relative bias** | | | | | | **Relative bias** | | | | | |
| $c$ | 0.020 | 0.061 | 0.13 | 0.10 | 0.15 | 0.14 | 0.043 | 0.083 | 0.17 | 0.053 | 0.034 | 0.13 |
| $K$ | 0.19 | 0.13 | 0.14 | 0.28 | 0.36 | 0.40 | 0.090 | 0.075 | 0.10 | 0.19 | 0.24 | 0.38 |
| $\theta$ | 0.040 | 0.055 | 0.055 | 0.022 | 0.043 | 0.039 | 0.028 | 0.027 | 0.12 | 0.057 | 0.032 | 0.074 |
| $\kappa$ | 0.0015 | 0.0019 | 0.00019 | $-0.0034$ | 0.0066 | 0.0065 | $-0.00030$ | 0.0062 | $-0.0020$ | 0.0041 | 0.00084 | $-0.0012$ |
| $\alpha$ | | N/A | | | N/A | | $-0.071$ | $-0.014$ | $-0.025$ | $-0.023$ | $-0.013$ | $-0.077$ |
| $\mu_a$ | | N/A | | | N/A | | 0.0077 | 0.0042 | 0.012 | $-0.0074$ | $-0.0020$ | 0.0087 |
| | **Relative RMSE** | | | | | | **Relative RMSE** | | | | | |
| $c$ | 0.43 | 0.43 | 0.59 | 0.57 | 0.62 | 0.62 | 0.35 | 0.39 | 0.57 | 0.49 | 0.51 | 0.60 |
| $K$ | 0.57 | 0.49 | 0.60 | 0.70 | 0.78 | 0.84 | 0.38 | 0.38 | 0.44 | 0.50 | 0.61 | 0.80 |
| $\theta$ | 0.30 | 0.29 | 0.33 | 0.28 | 0.28 | 0.30 | 0.30 | 0.32 | 0.45 | 0.29 | 0.30 | 0.32 |
| $\kappa$ | 0.060 | 0.049 | 0.042 | 0.073 | 0.075 | 0.073 | 0.041 | 0.035 | 0.031 | 0.045 | 0.042 | 0.043 |
| $\alpha$ | | N/A | | | N/A | | 0.50 | 0.096 | 0.039 | 0.86 | 0.20 | 0.11 |
| $\mu_a$ | | N/A | | | N/A | | 0.039 | 0.036 | 0.026 | 0.035 | 0.032 | 0.026 |

**Table S7. Coverage, relative bias, and relative RMSE of the posterior estimates for mutation model parameters and tree height, based on 100 replicate simulations, jointly inferring the genealogy and the model parameters under the misspecified Kingman coalescent model.** The table presents results for mutation model parameters and tree height. Seedbank-specific parameters without direct counterparts in the Kingman coalescent are excluded from inference. The synthetic datasets used here are identical to those in Tables 1. (A) Isochronous sampling (Scenario 1). The mutation rate was not inferred but held constant at either the active-state mutation rate $\mu_a$ (left) of the seedbank model or the tree-wide effective mutation rate $\mu_{\text{eff}}$ (right), computed as the edge-weighted average of $\rho_i$ (Eq. 3). (B) Serial sampling (Scenario 2). Because the Kingman coalescent does not distinguish between mutation rates of active and dormant states, a single mutation rate ($\mu$) is inferred and compared to the true $\mu_a$. For the case with $K = 1$ and $\alpha = 0.1$, results (marked "–") are excluded for comparisons with Table 1.

**(A) Isochronous sampling.**

| | $\mu = \mu_a$ | | | | | | $\mu = \mu_{\text{eff}}$ | | | | | |
|---|---|---|---|---|---|---|---|---|---|---|---|---|
| | $K = 1$ | | | $K = 10$ | | | $K = 1$ | | | $K = 10$ | | |
| | $\alpha = 0.1$ | $\alpha = 0.5$ | $\alpha = 0.99$ | $\alpha = 0.1$ | $\alpha = 0.5$ | $\alpha = 0.99$ | $\alpha = 0.1$ | $\alpha = 0.5$ | $\alpha = 0.99$ | $\alpha = 0.1$ | $\alpha = 0.5$ | $\alpha = 0.99$ |
| **Coverage** | | | | | | | | | | | | |
| $\kappa$ | 0.95 | 0.99 | 0.96 | 0.95 | 0.93 | 0.95 | 0.94 | 0.97 | 0.95 | 0.94 | 0.93 | 0.95 |
| $T_{\text{MRCA}}$ | 0.03 | 0.01 | 0.92 | 0.33 | 0.56 | 0.96 | 0.23 | 0.42 | 0.94 | 0.82 | 0.89 | 0.96 |
| **Relative bias** | | | | | | | | | | | | |
| $\kappa$ | 0.0074 | 0.00039 | $-0.0024$ | 0.0091 | 0.0015 | 0.0011 | 0.0071 | 0.00036 | $-0.0022$ | 0.0085 | 0.0018 | 0.0012 |
| $T_{\text{MRCA}}$ | $-0.43$ | $-0.26$ | $-0.0042$ | $-0.086$ | $-0.057$ | $-0.0027$ | $-0.087$ | $-0.038$ | 0.00047 | $-0.010$ | $-0.012$ | $-0.0019$ |
| **Relative RMSE** | | | | | | | | | | | | |
| $\kappa$ | 0.066 | 0.049 | 0.044 | 0.077 | 0.075 | 0.073 | 0.066 | 0.049 | 0.044 | 0.077 | 0.075 | 0.073 |
| $T_{\text{MRCA}}$ | 0.45 | 0.27 | 0.026 | 0.11 | 0.076 | 0.046 | 0.12 | 0.061 | 0.026 | 0.055 | 0.051 | 0.046 |

**(B) Serial sampling.**

| | $K = 1$ | | | $K = 10$ | | |
|---|---|---|---|---|---|---|
| | $\alpha = 0.1$ | $\alpha = 0.5$ | $\alpha = 0.99$ | $\alpha = 0.1$ | $\alpha = 0.5$ | $\alpha = 0.99$ |
| **Coverage** | | | | | | |
| $\kappa$ | – | 0.97 | 0.98 | 0.97 | 0.96 | 0.97 |
| $T_{\text{MRCA}}$ | – | 0.49 | 0.96 | 0.60 | 0.85 | 0.95 |
| $\mu$ | – | 0.0 | 0.96 | 0.12 | 0.35 | 0.97 |
| **Relative bias** | | | | | | |
| $\kappa$ | – | 0.0027 | $-0.0015$ | 0.0036 | 0.003 | 0.0011 |
| $T_{\text{MRCA}}$ | – | 0.0086 | 0.0014 | 0.00089 | 0.0016 | $-0.0023$ |
| $\mu$ | – | $-0.28$ | $-0.0043$ | $-0.097$ | $-0.055$ | 0.0043 |
| **Relative RMSE** | | | | | | |
| $\kappa$ | – | 0.033 | 0.029 | 0.043 | 0.043 | 0.043 |
| $T_{\text{MRCA}}$ | – | 0.067 | 0.024 | 0.029 | 0.020 | 0.016 |
| $\mu$ | – | 0.29 | 0.027 | 0.11 | 0.066 | 0.030 |

**Table S8. Coverage, relative bias, and relative RMSE of the posterior estimates for mutation model parameters and tree height, based on 100 replicate simulations inferred under the misspecified Kingman coalescent model, with either the mutation rate or the genealogy fixed (but not both).** (A) Serial sampling with a fixed mutation rate. The mutation rate was held constant at either the active-state mutation rate $\mu_a$ (left) of the seedbank model or the tree-wide effective mutation rate $\mu_{\text{eff}}$ (right), calculated as the edge-weighted average of $\rho_i$ (Eq. 3). (B) Serial sampling with a fixed genealogy. Model parameters are inferred given the true reduced (uncolored) genealogy. The synthetic datasets used here are identical to those in Tables 1.

**(A) Serial sampling, fixed mutation rate.**

**$\mu = \mu_a$**

| | $K = 1$ | | | $K = 10$ | | |
|---|---|---|---|---|---|---|
| | $\alpha = 0.1$ | $\alpha = 0.5$ | $\alpha = 0.99$ | $\alpha = 0.1$ | $\alpha = 0.5$ | $\alpha = 0.99$ |
| **Coverage** | | | | | | |
| $\kappa$ | – | 0.97 | 0.97 | 0.97 | 0.96 | 0.97 |
| $T_{\text{MRCA}}$ | – | 0.020 | 0.96 | 0.37 | 0.58 | 0.95 |
| **Relative bias** | | | | | | |
| $\kappa$ | – | 0.0057 | −0.0015 | 0.0041 | 0.0034 | 0.0010 |
| $T_{\text{MRCA}}$ | – | −0.18 | −0.0026 | −0.028 | −0.015 | −0.00094 |
| **Relative RMSE** | | | | | | |
| $\kappa$ | – | 0.033 | 0.029 | 0.043 | 0.043 | 0.043 |
| $T_{\text{MRCA}}$ | – | 0.19 | 0.014 | 0.037 | 0.022 | 0.013 |

**$\mu = \mu_{\text{eff}}$**

| | $K = 1$ | | | $K = 10$ | | |
|---|---|---|---|---|---|---|
| | $\alpha = 0.1$ | $\alpha = 0.5$ | $\alpha = 0.99$ | $\alpha = 0.1$ | $\alpha = 0.5$ | $\alpha = 0.99$ |
| **Coverage** | | | | | | |
| $\kappa$ | – | 0.97 | 0.97 | 0.97 | 0.96 | 0.96 |
| $T_{\text{MRCA}}$ | – | 0.45 | 0.97 | 0.73 | 0.88 | 0.95 |
| **Relative bias** | | | | | | |
| $\kappa$ | – | 0.0028 | −0.0015 | 0.0034 | 0.0031 | 0.0010 |
| $T_{\text{MRCA}}$ | – | −0.013 | 0.0012 | 0.000059 | 0.0017 | −0.00064 |
| **Relative RMSE** | | | | | | |
| $\kappa$ | – | 0.033 | 0.029 | 0.043 | 0.043 | 0.043 |
| $T_{\text{MRCA}}$ | – | 0.037 | 0.014 | 0.022 | 0.015 | 0.013 |

**(B) Serial sampling, fixed genealogy.**

| | $K = 1$ | | | $K = 10$ | | |
|---|---|---|---|---|---|---|
| | $\alpha = 0.1$ | $\alpha = 0.5$ | $\alpha = 0.99$ | $\alpha = 0.1$ | $\alpha = 0.5$ | $\alpha = 0.99$ |
| **Coverage** | | | | | | |
| $\kappa$ | – | 0.97 | 0.98 | 0.97 | 0.96 | 0.95 |
| $\mu$ | – | 0.0 | 0.96 | 0.0 | 0.09 | 0.94 |
| **Relative bias** | | | | | | |
| $\kappa$ | – | 0.0046 | −0.0015 | 0.0036 | 0.0032 | 0.0012 |
| $\mu$ | – | −0.26 | −0.0042 | −0.10 | −0.055 | −0.00017 |
| **Relative RMSE** | | | | | | |
| $\kappa$ | – | 0.033 | 0.029 | 0.043 | 0.043 | 0.043 |
| $\mu$ | – | 0.26 | 0.014 | 0.10 | 0.060 | 0.021 |

#### SI Figures

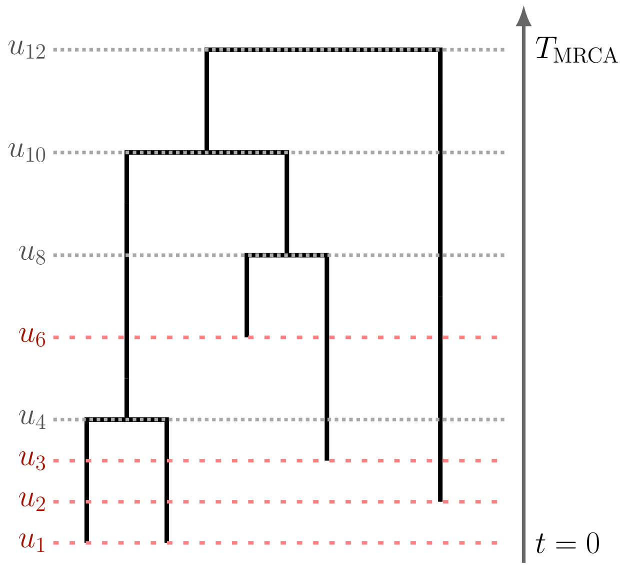


**Figure S2. Example of the reduced genealogy derived from the seedbank genealogy shown in Figure 1.**
